# Revealing architectural order with quantitative label-free imaging and deep learning

**DOI:** 10.1101/631101

**Authors:** Syuan-Ming Guo, Li-Hao Yeh, Jenny Folkesson, Ivan E. Ivanov, Anitha Priya Krishnan, Matthew G. Keefe, David Shin, Bryant Chhun, Nathan Cho, Manuel Leonetti, Tomasz J. Nowakowski, Shalin B. Mehta

## Abstract

Quantitative imaging of biological architecture with fluorescent labels is not as scalable as genomic or proteomic measurements. Here, we combine quantitative label-free imaging and deep neural networks for scalable analysis of complex structures. We reconstruct quantitative three-dimensional density, anisotropy, and orientation in live cells and tissue slices from polarization- and depth-resolved images. We report a computationally efficient variant of U-Net architecture that predicts a 3D fluorescent structure from its morphology and physical properties. We evaluate the performance of our models by predicting F-actin and nuclei in mouse kidney tissue. Further, we report label-free imaging of axon tracts and predict level of myelination in human brain tissue sections. We demonstrate the model’s ability to rescue inconsistent labeling. We anticipate that the proposed approach will enable quantitative analysis of architectural order across scales of organelles to tissues.

## Introduction

The function of living systems emerges from dynamic interaction of components over spatial and temporal scales that range many orders of magnitude. Although molecular components of biological systems can now be identified with scalable genomic and proteomic technologies, these technologies cannot capture dynamic interactions of components and are destructive measurements. Light microscopy is uniquely suited to study dynamic arrangement of molecules within the context of organelles, of organelles within the context of cells, and of cells within the context of tissues.

There is an outstanding need for scalable imaging technologies for mapping interactions among biological components. Widely used fluorescence microscopy methods report on labeled molecules within the crowded environment of biological system. However, it is difficult to visualize more than 7 components in a live sample (1) due to the stochastic nature of labeling, photo-toxicity, and the broad spectra of fluorescent proteins. Further, labeling of primary specimens and non-model organisms is still a challenge. In contrast, label-free imaging enables simultaneous and reproducible visualization of many biological structures with minimal photo-toxicity by measuring intrinsic physical properties of the sample. Label-free microscopy with phase contrast (2), differential interference contrast (DIC) (3), and polarization contrast (4, 5), has enabled discoveries of biological processes for almost a century.

Although label-free imaging is scalable, extracting quantitative measurements from label-free data is challenging due to two reasons: 1) in label-free images, physical properties of structures are encoded in complex modulation of the intensity, rather than the intensity magnitude, and 2) many structures seen in label-free data have similar physical properties and are therefore difficult to distinguish with classical analysis tools. To address these challenges, here we report a computational imaging method for joint measurement of density and structural anisotropy in 3D. We refer to density and anisotropy collectively as the architectural order in this paper. We also report computationally efficient deep neural networks that predict 3D fluorescent structures from 3D architectural order, thereby enabling analysis of specific structures within label-free data.

### Related work

Label-free imaging typically requires modifications of the optical path of a common microscope, as widely available cameras are sensitive only to the intensity (i.e., brightness) and wavelength (i.e., color) of light, not the relative times of arrival of light waves (i.e., phase) or the plane of oscillation of the electric field (i.e., polarization). Qualitative label-free imaging methods, such as phase contrast and DIC, transform the phase or polarization of light into intensity. Quantitative methods aim to reconstruct physical properties of the specimen using inverse algorithms based on image-formation models.

Phase microscopes report the optical path length ^1^, which is modulated by the specimen density. Optical path length and density can be measured from intensity modulation due to the propagation of light (7, 8), from interference between light scattered by the specimen with another beam whose phase is controlled (9–13), and from images of the specimen illuminated from multiple angles (14–16). Among these approaches, reconstruction of phase from propagation of light is experimentally the simplest approach, since the data can be acquired on any microscope with motorized focus drive. The weak object transfer function model of image formation (17–22) allows for robust reconstruction of phase from a stack of intensities measured in transmission.

Polarization microscopes measure retardance^1^, i.e., orientation-dependent optical path length, which is modulated by the structural anisotropy below the spatial diffraction limit and has been used to make measurements well below the resolution limit (23). Structural anisotropy can be measured using tunable polarization modulators in the illumination or the detection path of the microscope (24–28). Shribak *et al.* (13) developed a method that employed multiple liquid crystal modulators for joint analysis of phase and retardance in 2D, but not in 3D live specimens.

Quantitative phase imaging has been used to analyze diverse processes (9), such as, membrane mechanics, density of organelles, cell migration and more recently propagation of action potential (29). Polarized light imaging has enabled discovery of the dynamic microtubule spindle (5), assessment of structural integrity of meiotic spindles of oocytes in invitro fertilization (IVF) clinics (30), label-free imaging of white matter in adult human brain tissue slices (28, 31), and imaging of activity-dependent structural changes in brain tissue (32). It is apparent from above applications that joint imaging of density and anisotropy without label can enable new biological investigations. To our knowledge, approaches for combined measurement of density and anisotropy in live cells in 3D have not been reported.

Machine learning models have recently enabled identification of structures in diverse label-free images, opening new opportunities for scalable analysis of label-free data. Some examples include: 3D U-Net model for predicting multiple organelles in cells from brightfield and DIC images (33); inception model for in silico 2D labeling of nuclei, cell types, and cell state (34) from phase contrast and DIC images; generative adversarial models for 2D prediction of histopathological stains from quantitative phase (35) and auto-fluorescence (36); 3D U-Net model for segmentation of immunological synapse from diffraction tomography (37); random forest classifiers for recognizing carcinoma in colon tissue from Raman scattering (38). Among the above approaches, 3D U-Net models that translate label-free image stacks to a fluorescence stacks (33) are attractive, because they predict both localization and expression of specific molecules in 3D. However, learning anisotropic structures, such as cytoskeletal networks, can be challenging from brightfield data (33), because brightfield imaging is sensitive to density, but not structural anisotropy. In addition, density information in brightfield stacks is difficult to interpret and susceptible to imperfections in the light-path. Moreover, training and prediction using 3D U-Net is expensive - the computational cost required downsampling the data at the expense of spatial resolution (33).

There is an unmet need for computationally efficient 3D image translation models that can learn jointly from density and anisotropy measurements.

### Contributions

We recover density and anisotropy from polarization- and depth-resolved acquisition. Our measurements reveal structures across different biological scales, including lipid droplets, mitochondria, spindle, nucleoli, chromosomes, nuclei, nuclear envelope, plasma membrane, glomeruli and tubules in mouse kidney tissue, axon tracts in mouse and human brain tissue, and cell morphology in human brain tissue. The reconstruction relies on the description of polarization of light by a Stokes vector (39) and recovery of phase from 3D brightfield stack via deconvolution. Casting the image formation and reconstruction in Stokes formalism also provides an elegant representation of the microscope in terms of an instrument matrix (40–42), which enables robust calibration and background correction. Using the Stokes formalism and deconvolution algorithms reported here, we measure phase, retardance, slow axis ^2^, and degree of polarization of transparent specimen.

Next, we report a computationally efficient 2.5D U-Net architecture for translating 3D distribution of physical properties to fluorescence intensities. Our 2.5D architecture is related to architectures used to segment 3D MRI data from 2D projections of 3D data (43), but instead uses the information over the depth of field of the microscope. We chose to use information over the depth of field for prediction, because microscopes measure optical sections and not optical projections. Our 2.5D multi-channel models accurately translate brightfield, phase, retardance, and slow axis distributions over the depth of field of the microscope into fluorescence intensities in the corresponding 2D focal plane. 3D translation is achieved using the 2.5D model by predicting fluorescence at every slice of the output stack using neighboring slices of label-free stacks as inputs. Formulating the prediction problem over the depth of field allows us to work with substantially larger fields of view in a computationally efficient manner as compared to full 3D models. Using quantitative data as input removes artifacts and unintended correlations between label-free and fluorescence data, which improves accuracy of prediction and allows cleaner interpretation of predicted structure. Further, measured density and anisotropy can be interpreted jointly with predicted fluorescent structure to gain new insights in biology.

We applied our approach to mapping the axon tracts and myelination in a developing human brain across different ages. In contrast to brains of model systems (e.g., mouse and fruit fly), scalable mapping of the architecture of the human and other large brains poses significant challenges. For example, genetic tools are limited and not scalable for tracing the neuronal connectivity at high-throughput. To overcome this issue, we reasoned that joint imaging of density and structural anisotropy will allow us to more thoroughly describe the structures of human brain tissue.

Collectively, above contributions establish a novel approach for imaging architectural order in living systems across biological scales and analyzing it with a judicious combination of physics-driven and data-driven modeling approaches.

## Results

### Reconstructing density and anisotropy

Simultaneous recovery of multiple properties requires a model of image formation accurate enough to account for partial polarization of light. Previous reconstruction algorithms (25, 26) for transmission polarized light microscopy employed Jones calculus (39, Ch.10) to describe image formation, which assumes fully polarized illumination. However, LEDs or lamps used for transmission microscopy lead to partially polarized illumination that cannot be accounted for by Jones calculus. Stokes vector representation of light and Mueller matrix representation of the optical components cleanly capture the full state of polarization of light (39, 41, 44), including partial polarization. We previously developed Stokes representation of fluorescence polarization for simultaneous recovery of concentration, alignment, and orientation of fluorophores imaged with instantaneous fluorescence polarization microscope (42). Fluorescence polarization microscopy also gives rise to partially polarized emission, since independent emission events integrated through the imaging aperture are mutually incoherent. In this paper, we employ the Mueller calculus to describe transmission imaging with partial polarization. We exploit the same framework for calibration, background correction, and reconstruction of physical properties.

We implemented automated polarization-resolved and depth-resolved imaging (fig. 1A, methods). We used 5 elliptical polarization states for sensitive detection of specimen retardance (25, 26)(methods). Our image formation model represents the specimen’s physical properties in terms of Mueller coefficients. We recover background-corrected Mueller coefficients (methods) of the specimen at every voxel(fig. 1B). We reconstruct brightfield, phase, retardance, slow axis, and degree of polarization stacks from stacks of Mueller coefficients (fig. 1C, methods). We developed a two-step background correction method that enables detection of low anisotropy of the biological structures even in the presence of high, non-uniform background resulting from the optics or imaging chamber (methods,fig. 2-supplement S4).

**Fig. 1.**
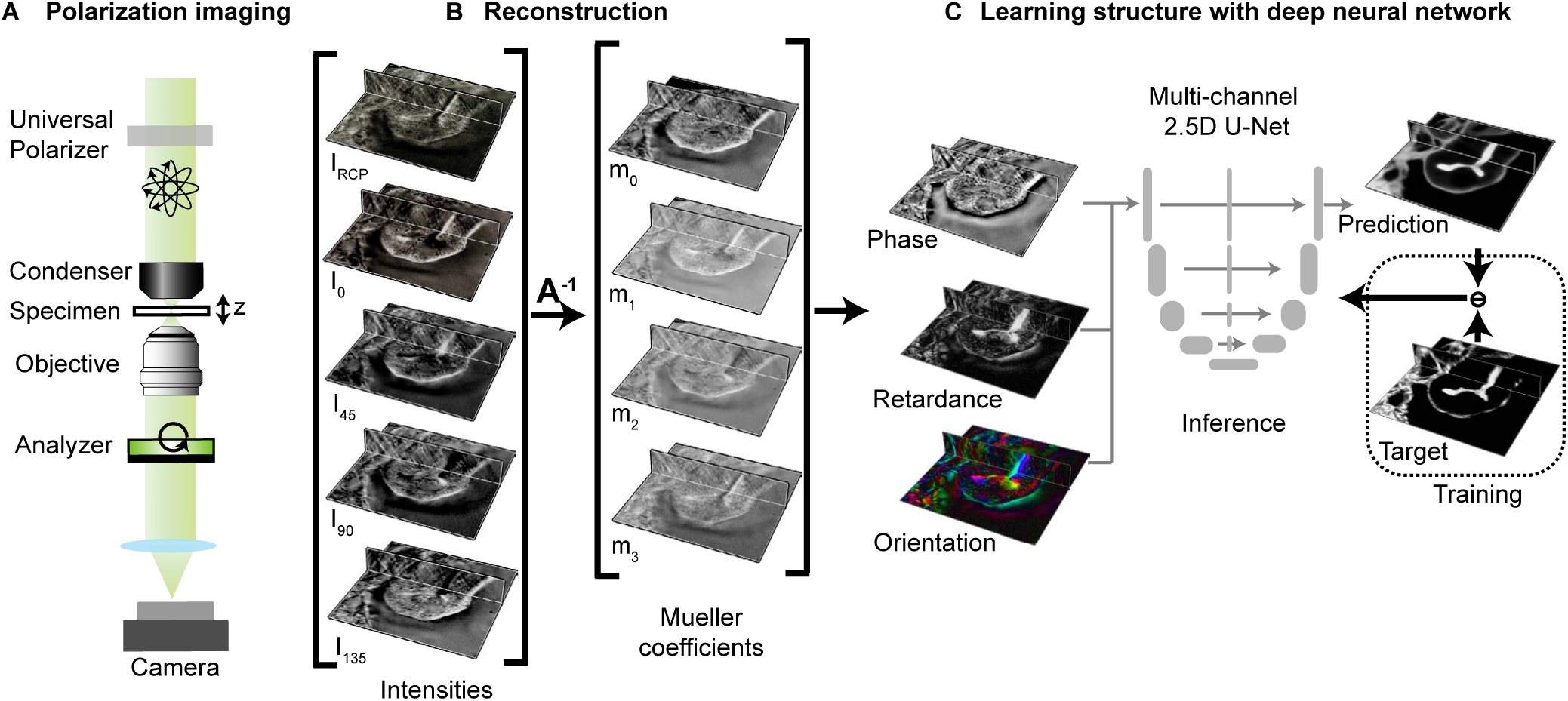
Label-free prediction of a structure from density and anisotropy measurements: **(A)** Light path of label-free polarization-resolved and depth-resolved imaging microscope. The measured intensities encode spatial distributions of refractive index and birefringence of the specimen. Polarization states are generated by a calibrated liquid-crystal universal polarizer that can be controlled electronically. **(B)** Mueller coefficients of the specimen are computed for every voxel from polarization-resolved intensities using the inverse of instrument matrix **A**^*−*1^ (see eq. (9)). Mueller coefficients retrieved from the background region are used to correct imperfections in the light path (fig. 2-supplement S4). Assuming that the specimen is transparent, complementary physical properties of the specimen are reconstructed from Mueller coefficients: phase, retardance, orientation of slow axis, brightfield (not shown), and degree of polarization (not shown). **(C)** Multi-channel, 2.5D U-Net model is trained to predict 3D fluorescent structures from label-free measurements. During training, pairs of label-free images and fluorescence images are supplied as inputs and targets to the U-Net model. The model is optimized by minimizing the difference between the model prediction and the target. During inference, only label-free images are supplied as input to the trained model to predict 3D fluorescence images.

**Fig. 2.**
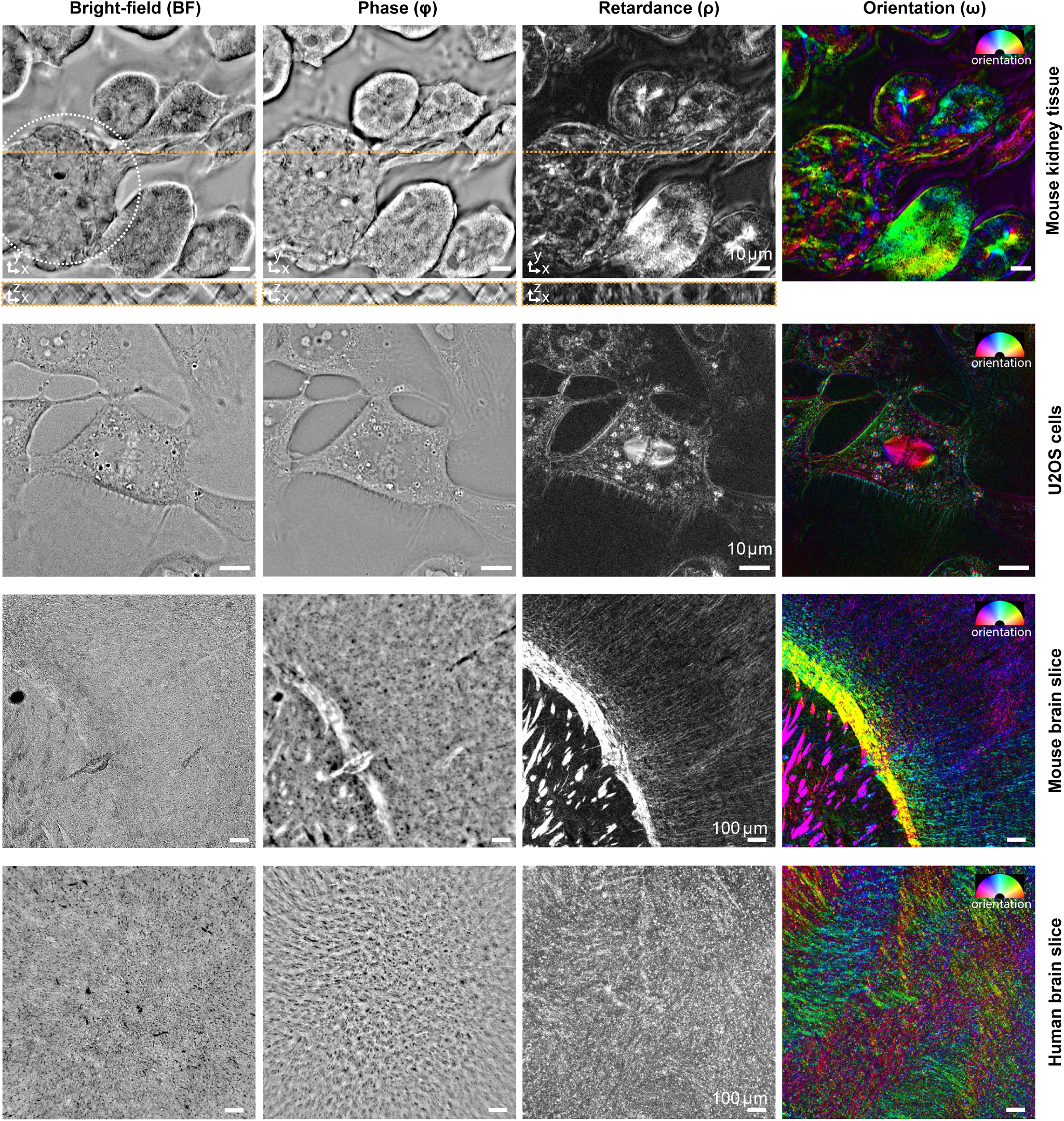
Complementary contrasts of phase and retardance report diverse biological structures: Brightfield (BF), Phase (*φ*), retardance (*ρ*), and orientation (*ω*) images of mouse kidney tissue, mammalian cell line (U2OS), mouse brain tissue section, and human brain tissue section are shown. The orientation image represents slow axis of the specimen with color (hue) and retardance with brightness. In kidney tissue, nuclei appear as smooth patches of low density and low retardance, while tubules stand-out in retardance. Glomelurus, a network of small blood vessels, is identified by circular outline in the brightfield image of kidney tissue. In U2OS cells, chromatin, lipid droplets, membranous organelles, and cell boundaries are visible due to variations in density, while microtubule spindle, lipid droplets, and cell boundaries are visible due to their ordered structure. In mouse brain slices, axon tracts are more visible in phase, retardance, and orientation images compared to brightfield images, with slow axis perpendicular to the direction of the bundles. Similar contrast improvement was observed in developing human brain tissue slice with less ordered tracts due to the early age of the donor. 3D stacks of mouse kidney tissue and U2OS cell were acquired with 63x 1.47 NA oil objective and 0.9 NA illumination, whereas images of mouse and human brain tissue were acquired with 10x 0.3 NA air objective and 0.2 NA illumination.

Figure 2 shows background-corrected images of a mouse kidney tissue slice, U2OS cells (bone cancer cell line), mouse brain slice, and developing human brain slice in brightfield, phase, retardance, and orientation contrasts. Both the phase and brightfield images report density variation in the specimen. However, in brightfield images, optically dense structures appear in brighter contrast than the background on one side of the focus, almost no contrast at the focus, and darker contrast than the background on the other side of the focus (note the contrast variations in images of nuclei in the XZ section of the brightfield image of the mouse kidney tissue and in Video 1). This contrast variation along the optical axis, described by the transport of intensity equation (7), makes proper interpretation of density from brightfield images challenging. We developed a phase reconstruction algorithm (methods) that translates this axial contrast modulation into a more quantitative contrast that is proportional to the density, i.e. denser structures consistently appear in brighter contrast than the background in the image. The retardance image is proportional to the degree of orientational order among molecules. The slow axis orientation image reports the orientation in which the specimen is the densest. In mouse kidney tissue, the retardance image highlights capillaries within glomeruli, and brush borders in convoluted tubules, among other components of the tissue. The nuclei appear in darker contrast in the retardance image because of the relatively less ordered structure of chromatin.

In the dividing U2OS cell (Video 2), the phase image shows contrast consistent with density of various cellular structures including cytoplasm, lipid vesicles, nucleoli, and chromosomes compared to the brightfield image where the contrast of the structures varies depending on their location relative to the focus. Similarly, in mouse and developing human brain tissue sections, the phase image identifies axon tracts better than brightfield image, because of variations in their density (fig. 2-supplement S1). The background corrected retardance and orientation in U2OS cells (Video 2) show dynamics of membrane boundaries, spindle, and lipid droplets. We note that the two-step background correction (methods) is essential to remove biases in the retardance and orientation images, but not for phase image (fig. 2-supplement S4). The retardance and orientation images of mouse and human brain slices in fig. 2 distinctly report on axon tracts. The birefringence of the axons arises from neurofilaments that have higher density along the axon axis and myelin sheath that has higher density perpendicular to the axon axis (45). Due to the high birefringence of myelinated axons in mouse brain slice, we see a slow axis perpendicular to the direction of the axon tracts. Figure 2-supplement S2 shows stitched retardance and orientation images of a whole mouse brain slice, in which not just the white matter tracts, but also changes in the orientation of axons across different cortical layers are visible.

We show degree of polarization measurements in fig. 2-supplement S3. It is worth clarifying the difference between retardance images shown in fig. 2 and degree of polarization images. The retardance variations arise from single scattering events within the specimen that alter the polarization, but do not reduce the degree of polarization. The degree of polarization on the other hand reports multiple scattering events that reduce the polarization of light. In the future, we plan to pursue models that account for diffraction and scattering effects in polarized light microscopy and enable more precise retrieval of specimen properties.

Data shown in fig. 2 and in Video 3 report simultaneous, quantitative measurements of density, structural anisotropy, and orientation in 3D biological specimens, for the first time to our knowledge. In the next sections, we discuss how these complementary label-free measurements enable prediction of fluorescence images of different types of structures. The Python code for reconstruction is available at https://github.com/mehta-lab/reconstruct-order.

### Computationally efficient 3D prediction using 2.5D residual U-Net

Joint optimization of optical contrast, architecture of the deep neural network, and the training process is key to successful analysis of structures of interest. Label-free measurement of density and anisotropy simultaneously visualize several structures that are normally imaged with fluorescence labeling. To enable automated analysis of localization and expression of specific molecules, we sought to develop deep convolutional neural network models that translate 3D label-free stacks into 3D fluorescence stacks. We optimized our model architecture using mouse kidney tissue sections where F-actin and nuclei are labeled.

We adapted the U-Net architecture that has been widely successful in image segmentation and translation tasks (33, 37, 46, 47). U-Net’s success arises from its ability to exploit image features at multiple spatial scales. It uses skip connections between the encoding and decoding blocks that give decoding blocks access to low-complexity, high-resolution features in the encoding blocks. We added a residual connection between the input and output of each block to speed up training of the model (47, 48). Prior work (33) on predicting fluorescence stacks from brightfield stacks has shown that 2D models result in discontinuous predictions along the z-axis as compared to 3D translation models. However, training and prediction with 3D translation models are computation and memory intensive. Further, typical microscopy stacks are bigger in their extent in the focal plane (~2000×2000 pixels) and smaller in extent along the optical axis (usually *<* 40 Z slices). But 3D U-Net model requires sufficiently large number of Z slices (*>*= 64) to extract meaningful high level features in Z-dimension as the input is isotropically down-sampled in the encoding path of the U-Net. Therefore, use of 3D translation models often requires upsampling the data in Z, which increases data size and makes training 3D translation model more computationally expensive.

To reduce the computational cost while maintaining the optical resolution of the data and high accuracy of prediction, we evaluated the prediction accuracy as a function of model dimensions for a highly ordered structure (F-actin) and for less ordered structure (nuclei) in mouse kidney tissue. We evaluated three model architectures to predict fluorescence volumes: slice→slice (2D in short) models that predict 2D fluorescence slices from corresponding 2D label-free slices, stack→slice (2.5D in short) models that predict the central 2D fluorescence slices from a stack of neighbor label-free slices, and stack→stack (3D in short) models that predict 3D fluorescent stacks from label-free stacks. For 2D and 2.5D models, 3D translation is achieved by using the model to predict fluorescence in each plane of the stack. See methods and fig. 3-supplement S1 for the description of the network architecture and training process. We used Pearson correlation coefficient and structural similarity index (SSIM) (49) between predicted fluorescent stacks and target fluorescent stacks from the test fields of view to evaluate the performance of the models (methods). We report these metrics on the test set (table 1, table 2, table 3), which was not used during the training.

**Table 1.** Accuracy of 3D prediction of F-actin from retardance stack using different neural networks: Above table lists median values of the Pearson correlation (*r*) and structural similarity index (SSIM) between prediction and ground truth F-actin volumes. We report accuracy metrics for Slice*→*Slice (2D), Stack*→*Slice (2.5D), and Stack*→*Stack (3D) models trained to predict F-actin from retardance using Mean Absolute Error (MAE or L1) loss. We segmented target images with Rosin threshold to discard tiles that mostly contained background pixels. To dissect the differences in prediction accuracy along and perpendicular to the focal plane, we computed (methods) test metrics separately over XY slices (*r*_*xy*_, SSIML_*xy*_) and XZ slices (*r*_*xz*_, SSIM_*xz*_) of the test volumes, as well as over entire test volumes (*r*_*xyz*_, SSIM_*xyz*_).

| Translation model | Input(s) | $r_{xy}$ | $r_{xz}$ | $r_{xyz}$ | SSIM<sub>xy</sub> | SSIM<sub>xz</sub> | SSIM<sub>xyz</sub> |
| --- | --- | --- | --- | --- | --- | --- | --- |
| Slice-> Slice (2D) | $\rho$ | 0.82 | 0.79 | 0.83 | 0.78 | 0.71 | 0.78 |
| Stack-> Slice (2.5D, $z = 3$) | $\rho$ | 0.85 | 0.83 | 0.86 | 0.80 | 0.75 | 0.81 |
| Stack-> Slice (2.5D, $z = 5$) | $\rho$ | 0.86 | 0.84 | **0.87** | 0.81 | 0.76 | 0.82 |
| Stack-> Slice (2.5D, $z = 7$) | $\rho$ | **0.87** | **0.85** | **0.87** | **0.82** | **0.77** | 0.83 |
| Stack-> Stack (3D, $z = 96$) | $\rho$ | 0.86 | 0.84 | 0.86 | **0.82** | 0.76 | **0.85** |

**Table 2.** Accuracy of prediction of F-actin in mouse kidney tissue as a function of input channels: Above table lists median values of the Pearson correlation (*r*) and structural similarity index (SSIM) between prediction and target volumes of F-actin. We evaluated combinations of brightfield (BF), phase (*φ*), retardance (*ρ*), orientation x (*ω*_*x*_), and orientation y (*ω*_*y*_), as input. Model training conditions and computation of test metrics is described in table 1.

| Translation model | Input(s) | $r_{xy}$ | $r_{xz}$ | $r_{xyz}$ | SSIM<sub>xy</sub> | SSIM<sub>xz</sub> | SSIM<sub>xyz</sub> |
| --- | --- | --- | --- | --- | --- | --- | --- |
| Stack-> Slice (2.5D, $z = 5$) | $\rho$ | 0.86 | 0.84 | 0.87 | 0.81 | 0.76 | 0.82 |
|  | BF | 0.86 | 0.84 | 0.86 | 0.82 | 0.77 | 0.83 |
| | $\phi$ | 0.87 | 0.85 | 0.88 | **0.83** | 0.78 | 0.84 |
| | $\phi, \rho, \omega_x, \omega_y$ | **0.88** | **0.87** | **0.89** | **0.83** | **0.80** | **0.85** |
| | BF, $\rho, \omega_x, \omega_y$ | **0.88** | **0.87** | **0.89** | **0.83** | 0.79 | **0.85** |

**Table 3.**
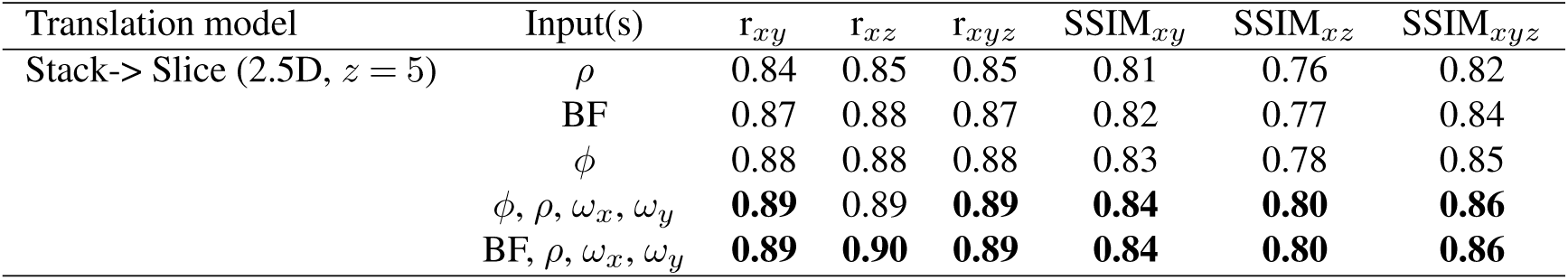
Accuracy of prediction of nuclei in mouse kidney tissue: Above table lists median values of the Pearson correlation (*r*) and structural similarity index (SSIM) between prediction and target volumes of nuclei. See table 2 for description.

**Fig. 3.**
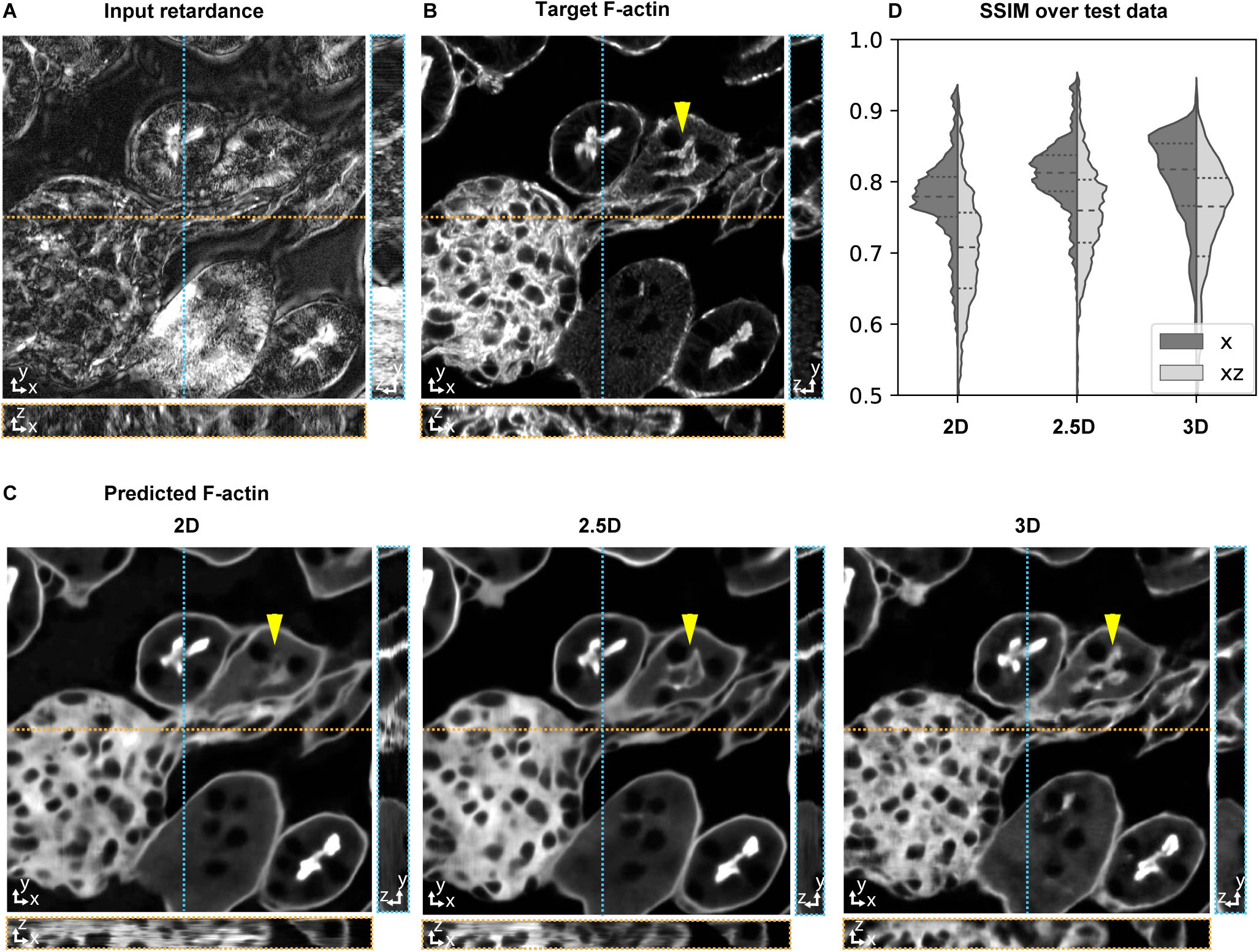
Accurate 3D prediction of fluorescent structure with 2.5D U-Net: Orthogonal sections (XY - top, XZ - bottom, YZ - right) of 3D volumeric test images of (A) retardance, (B) Experimental fluorescence of F-actin stain (target image), and (C) F-actin predicted from the retardance image using different U-Net architectures. (D) Violin plots of structral-similarty metric (SSIM) between images of predicted and experimental stain in XY and XZ planes. The horizontal dashed lines in the violin plots indicate 25^th^ quartile, median, and 75^th^ quartile of SSIM. The yellow triangle in C highlights a tubule structure, whose prediction can be seen to improve as the model has access to more information along Z. The same field-of-view is shown in fig. 2, Video 1, Video 4, and Video 5.

Figure 3A and fig. 3B show orthogonal slices of retardance and experimental F-actin stacks from the test set around a glomerulus and surrounding tissue, while fig. 3C shows orthogonal slices through the F-actin volumes predicted using 2D, 2.5D, and 3D models trained on retardance as the input. Glomeruli are complex multi-cellular, three-dimensional structures in kidney that perform filtration (50). The predictions with 2D models show discontinuity artifacts in the structure along the depth (fig. 3C, Video 4), as also observed in prior work (33). The 2.5D model predicts smoother structures along the Z dimension and improves the fidelity of F-actin prediction in the XY plane, (fig. 3C, table 1), with higher prediction accuracy as the number of z-slices in the 2.5D model input increases. The 3D model further improves the fidelity and continuity prediction along the depth (fig. 3C and Video 4). The distribution of SSIM along XY and XZ slice (fig. 3D, table 1) shows that the 2.5D model approaches the prediction accuracy of the 3D model, which is further improved when using complementary label-free properties as illustrated in the next section.

We note that we could train 2.5D model with ~3× more learnable parameters than 3D model (methods) while using fewer computational and memory resources. In our experiments, training a 3D model with 1.5*M* parameters required 3.2 days, training a 2D model with 2*M* parameters required 6 hrs, and training a 2.5D model with 4.8*M* parameters and 5 input z-slices required 2 days, using *∼* 100 training volumes.

The Python code for training our variants of image translation models is available at https://github.com/czbiohub/microDL.

### Predicting structures from multiple label-free contrasts improves accuracy

Considering the trade-off between computation speed and model performance, we adopted 2.5D models with 5 input Z-slices to explore how combinations of label-free inputs affect the accuracy of prediction of fluorescent structures.

We found that when multiple label-free measurements are jointly used as inputs, both F-actin and nuclei are predicted with higher fidelity compared to when only a single label-free measurement is used as the input (table 2 and table 3). Figure 4 A-C shows structural differences in the predictions of the same glomerulus as fig. 3. These observations from a representative field of view generalize to the entire test set as illustrated by the distribution of SSIM between slices of ground-truth and predictions in fig. 4C. The continuity of prediction along Z-axis improves as more label-free contrasts are used for prediction (Video 5). These results indicate that our model leverages information in complementary physical properties to predict target structures. We note that using complementary label-free contrasts boosted the performance of 2.5D models to exceed the performance of 3D single-channel models without significantly increasing the computation cost (compare table 1 and table 2). The prediction accuracy for fine F-actin structures also improves when complementary contrasts are used as input (Video 4 and Video 5).

**Fig. 4.**
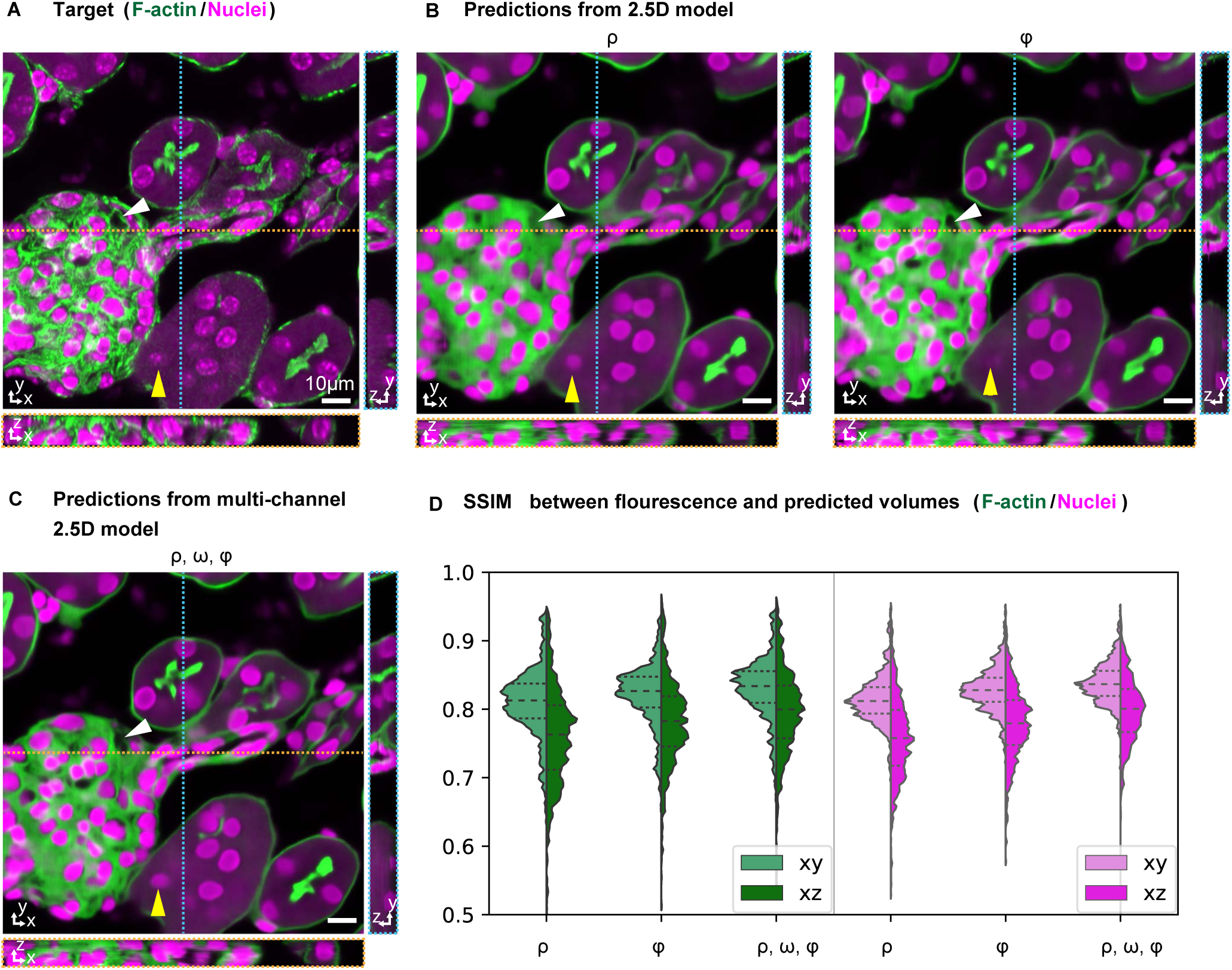
Prediction accuracy improves with multiple label-free contrasts as inputs: 3D predictions of ordered F-actin and nuclei from different combinations of label-free contrasts using 2.5D U-Net model. **(A)** Fluorescent stain for a field of view from the test set. Phalloidin-labeled F-actin in shown green and DAPI labeled nuclei is shown in magenta. **(B)** F-actin and nuclei distribution predicted with models trained on retardance (*ρ*) and phase (*φ*) alone are shown. **(C)** F-actin and nuclei distribution predicted with models trained with combined input of retardance, orientation, and phase. **(D)** Violin plots of structral-similarty metric (SSIM) between images of predicted and experimental stain in XY and XZ planes. The horizontal dashed lines in the violin plots indicate 25^th^ quartile, median, and 75^th^ quartile of SSIM. The yellow triangle and white triangle point out structures missing in predicted F-actin and nuclei distributions when only one channel is used as an input, but predicted when all channels are used. Label-free inputs used for prediction are shown in fig. 2 and Video 1.

Interestingly, when only a single contrast is provided as the input, a model trained on phase images has higher prediction accuracy than the model trained on brightfield images. This is possibly because the phase image has consistent, quantitative contrast along z-axis, while the depth-dependent contrast in brightfield images makes the learning task more challenging. This improvement of using phase over bright-field images, however, is not observed when the retardance and orientation images are included as inputs as well, possibly because the quantitative contrast in retardance and orientation images provides sufficient information for prediction of nuclei. We note that fine details of striated F-actin and nucleoli are missing even in the multi-contrast 2.5D model prediction (Video 5). This may be due to limited information content in the input images or due to the model not being trained on sufficient examples of fine features.

In conclusion, above results show that 2.5D multi-contrast models predict 3D structures with as high an accuracy as 3D U-Net models, but have multiple practical advantages that facilitate scaling of the approach. In addition, the results show that structures of varying density and order can be learned with higher accuracy when complementary physical properties are used as inputs.

### Mapping axon tracts and their myelination in sections of developing human brain

We next evaluate the possibility of mapping axon tract orientation and myelination in a postmortem human fetal brain without label. Studies in prenatal (51) and postnatal (52) human brains suggest that human brains develop and are wired differently as compared to model systems such as mice. Development of myelination is essential for long-range neural transmission and, consequently, emergence of coordination among different parts of brain (52). Scalable imaging of myelination and connectivity in developing human brain has the potential to enable studies that address outstanding questions of fundamental importance and clinical relevance. Label-free imaging of white matter is already being pursued in adult human brains (28) with polarized light. However, mapping myelination and axon tract orientation during human development requires higher sensitivity to changes in density and anisotropy, since contrast is much lower than developed human brain. Deep learning models that translate this data into quantifiable maps of myelination can facilitate analysis of the complex information contained in such data.

We imaged brain sections from two different ages, gestational week (GW) 24 (fig. 5A–E, fig. 5-supplement S1) and GW20 (fig. 5F–J, fig. 5-supplement S2), which correspond to the earliest stages of oligodendrocyte maturation and early myelination in the cerebral cortex (51). The stitched retardance and orientation images show the orientation of the axon tracts in different regions of the brain that are not accessible with brightfield or phase imaging, with slow axis orientation perpendicular to the direction of the axon tracts, similar to the observations on the mouse brain section (compare with fig. 2-supplement S2). The retardance of the white matter in subplate is higher than cortical plates in both time points, which is consistent with the reduced myelin density in the cortical plate relative to the white matter. Importantly, thanks to the calibration and background correction methods (methods), we are able to detect the orientation of axons in cortical plates even though myelin density is low early during development.

**Fig. 5.**
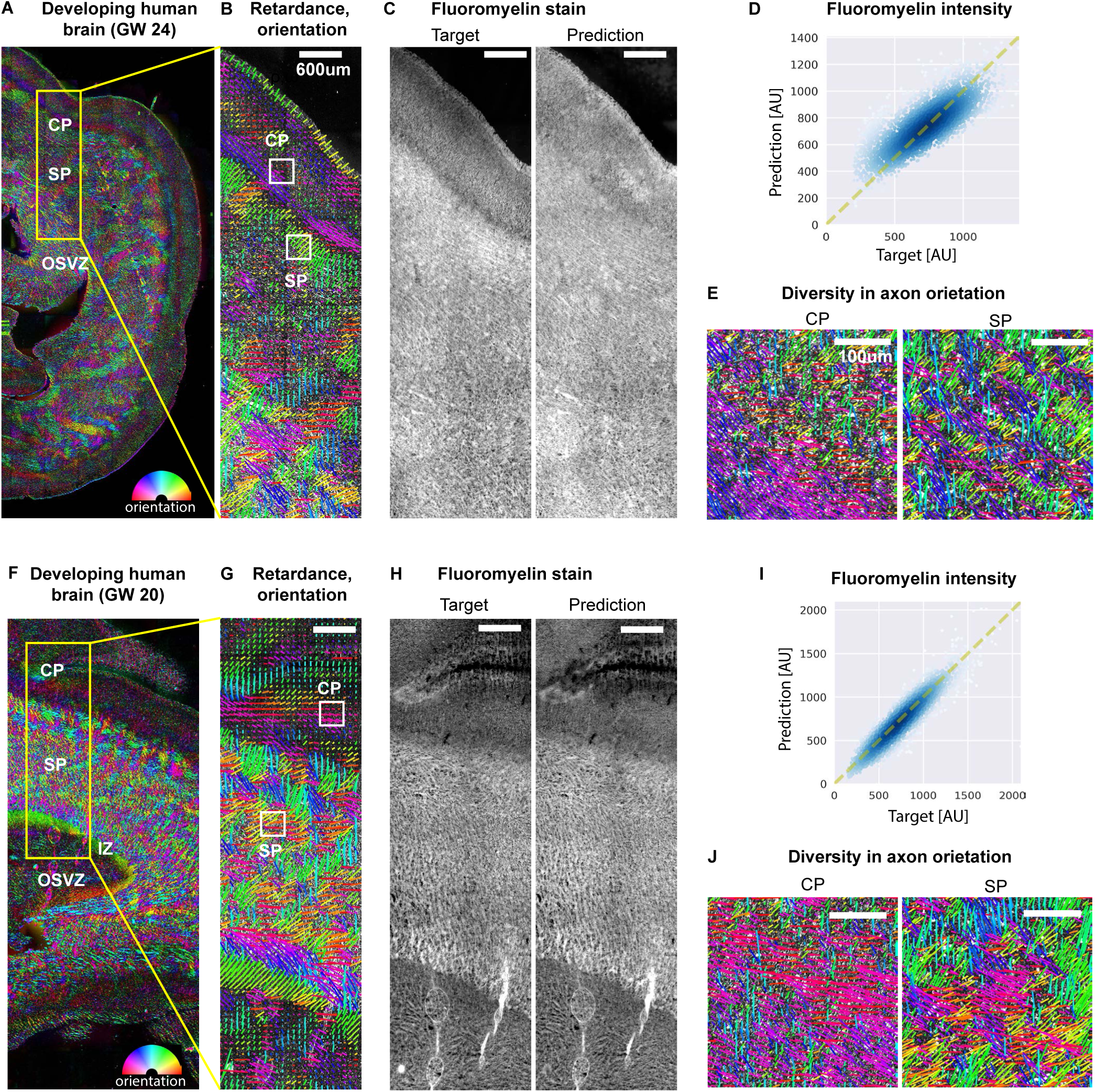
Label-free mapping of myelination and axon tracts in developing human brain tissue section: **(A)** Stitched image of retardance and orientation of a gestational week 24 (GW24) brain section from the test set. Different regions of the brain can be identified in the label-free images, CP: cortical plate; SP: subplate; OSVZ: outer subventricular zone. The orientation is encoded by color according to legend shown in bottom-right. **(B)** Region of interest over cortical plate and subplate showing retardance and orientation as lines. The orientation is represented by both colors and orientations of the lines, and retardance is represented by length of the lines. **(C)** Experimental FluoroMyelin stain (target) and the predicted fluoromyelin stain from the label-free contrasts with the most accurate model identified in table 4. **(D)** Scatter plot of target and predicted intensity. Yellow dashed line indicates the function y=x. **(E)** Insets from **(B)** of retardance and orientation highlight difference in pattern of orientation of tracts in cortical plate and mixed orientation of tracts in subplate. **(F – J)** Same as (A–E), but for GW20 sample. IZ: intermediate zone.

In contrast to predictions reported in fig. 3 and fig. 4, here we performed 2D instead of 3D phase reconstruction for human brain slice dataset as the archival tissue (12 *µ*m thick) was thinner than the depth of field (~16 *µ*m) of the low magnification objective (10X) we used for imaging large areas. The stitched 2D reconstructed phase shows tissue morphology and axon tracts with significantly higher contrast when compared to the brightfield image (fig. 5-supplement S1A and fig. 5-supplement S2A). This is because 2D phase reconstruction is more sensitive to density variation integrated along the z-axis compared to the brightfield imaging.

**Table 4.**
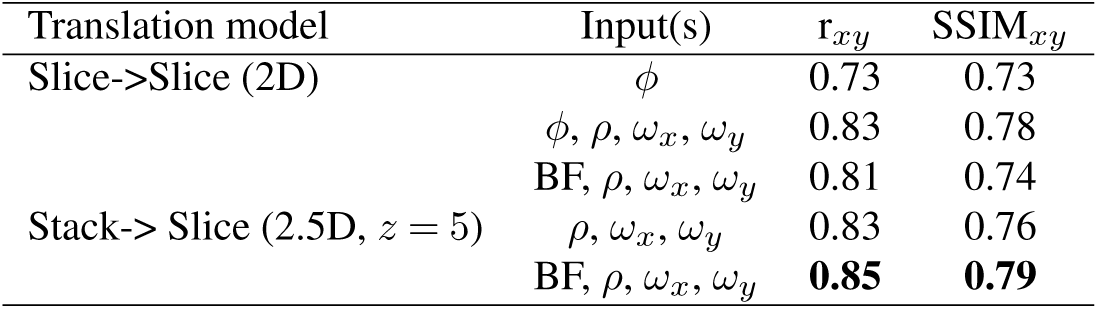
Accuracy of prediction of FluoroMyelin in human brain tissue slices across two developmental points (GW20 and GW24): Above table lists median values of the Pearson correlation (*r*) and structural similarity index (SSIM) between predictions of image translation models and target fluorescence. We evaluated combinations of retardance (*ρ*), orientation x (*ω*_*x*_), orientation y (*ω*_*y*_), phase (*φ*), and brightfield (*BF*) as inputs. These metrics are computed over 15% of the fields of view from two GW20 datasets and two GW24 datasets that were not used during training. The 2D models take *∼* 4 hours to converge, whereas 2.5D models take *∼* 64 hours to converge.

Interestingly, the pattern of orientation in cortical plate at GW24 (fig. 5E) suggests layered arrangement of neurons, while the pattern of orientation in subplate at GW24 shows criss-crossing arrangement of neurons. These differences between orientation pattern in cortical plate and subplate are less pronounced in data acquired from GW20 tissue (fig. 5I). To our knowledge, above data is the first report of label-free imaging of myelination and axon tract orientation in prenatal brain tissue. These data illustrate that the sensitivity and resolution of our approach

### Predicting myelination in sections of developing human brain

We explored how information in the label-free measurements can be used to predict myelination, which serves as a proxy for development of white matter. We applied our image translation approach to the task of predicting myelination in the developing human brain at 2 different ages. Myelination is visualized using lipophilic organic dye FluoroMyelin that preferentially binds to myelin (53).

To test how well our models generalize to different brain sections not seen by the model during the training stage, we ran model inference on images of a GW24 and GW20 sections that were not included in training, validation, or test set (results shown in fig. 5 and its supplements). The models were able to predict the axon tracts and myelination in these sections well, with increasing accuracy as we included more label-free channels as the input. Proper data normalization was essential for predicting the intensity correctly across different fields-of-view in large stitched images. Due to the batch variation of the staining and imaging process, we found that normalizing the images over whole the dataset gives the most accurate intensity prediction (fig. 5-supplement S3).

We trained multi-contrast 2D and 2.5D models with different combinations of label-free input contrasts and FluoroMyelin as the target to predict. To improve the model accuracy and generalize the model prediction to different development ages and different types of sections of the brain, we pooled imaging datasets from GW20 and GW24, with 2 different brain sections for each age. The pooled dataset was then split into training, validation, and test set. Similar to the observations on the mouse kidney tissue dataset, 2.5D model with brightfield, retardance, and orientation as the input has the highest SSIM and Pearson correlation over the test set (table 4). The linearity between predicted and target FluoroMyelin with scatter plots (fig. 5D and fig. 5I) shows that our model reliably translates the complex information in density and anisotropy to expression of FluoroMyelin.

Notably, the 2D model with phase, retardance, and orientation as the input has scores close to the best 2.5D model but the training takes 3.7 hrs to converge, while the best 2.5D model takes 64.7 hrs to converge (table 4). This is most likely because the 2D phase reconstruction captures the density variation encoded in the brightfield Z-stack that is informative for the model to predict axon tracts accurately.

#### Rescue of inconsistent label

Another useful feature of our image translation approach is that prediction results are robust to experimental variation in fluorescent stains. For example, we found that antifade chemical in the mounting media quenched the fluorescence of FluoroMyelin and created dark patches in FluoroMyelin (fig. 6) images. However, the quenching of dye had no impact on the physical properties we measure in the label-free channels. Therefore, the model trained on images without artifacts from fluorescence quenching predicted the expected staining pattern even where experimental stain failed. This robustness is particularly valuable for precious tissue specimens such as archival prenatal human brain tissue.

**Fig. 6.**
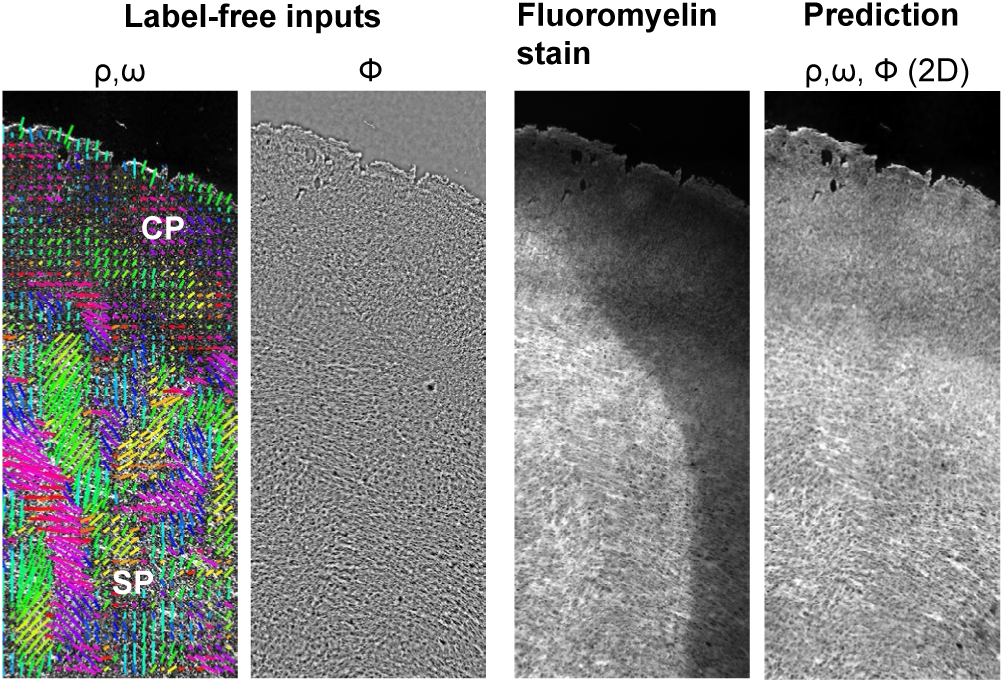
Model predicted stain is robust to experimental variation in staining: ROI of GW24 brain section shows artifacts in FluoroMyelin stain due to quenching of fluorescence by mounting medium. FluoroMyelin stain predicted by the model does not show the artifacts observed in experimental FluoroMyelin stain. (Left to right) retardance, orientation, and phase input of the 2D model; experimental FluoroMyelin stain; predicted stain from multi-contrast 2D model. CP: cortical plate; SP: subplate.

## Discussion

We have reported an innovative approach for label-free measurement of density and anisotropy from 3D polarization-resolved acquisition. We discuss how we elected to balance the trade-offs and the future directions of research.

The phase and degree of polarization information is inherently present in polarization-resolved acquisition, but can now be reconstructed thanks to our accurate forward models and corresponding reconstruction algorithms. As compared to joint imaging of density and anisotropy with orientation-independent differential interference contrast and orientation-independent PolScope (13), our method uses only one polarization modulator that simplifies calibration. However, our simpler light path achieves lower depth sectioning compared to differential interference contrast. Our method provides diffraction-limited imaging of density and ansiotropy in live cells, as evident from the 3D movie of organelles (fig. 2, Video 2, Video 3). Our open-source Python software is free to use for non-profit research. We anticipate the modularity of the optical path and the availability of software to facilitate adoption.

Our approach of recovering phase from propagation of light, reports the local phase variation and not the absolute phase. Measurement of absolute phase would require use of interference with a reference beam (10, 11). Nonetheless, we can still visualize many biological processes by using the relative phase variation. Further, we employ partially coherent illumination, i.e., simultaneous illumination from multiple angles. Partially coherent illumination improves spatial resolution, depth sectioning, and robustness to imperfections in the light path away from the focal plane.

We also note that, similar to other present polarization-resolved imaging systems (28, 42), our approach reports projection of the anisotropy onto the focal plane. Anisotropic structures, such as axon bundles, appear isotropic to the imaging system when they are aligned along the optical axis of the imaging path. Recovering true 3D anisotropy along with 3D density using forward models that account for diffraction effects in the propagation of polarized light is the an important open area of research.

Polarization-sensitive imaging has also been performed in reflection mode, most commonly with polarization sensitive optical coherence tomography (PS-OCT). PS-OCT has been used to measure round-trip birefringence and diattenuaton of diverse tissues, e.g., of brain tissue (54). But determination of the material axes in the reflection mode is confounded by the fact that light passes through the specimen in two directions. The reconstruction and background correction algorithms in PS-OCT primarily rely on Jones calculus, since OCT is a coherent interferometer and intensity recorded in individual speckle is fully polarized (55). However, PS-OCT practitioners employ degree of polarization uniformity (55) over several speckles to analyze depolarization due to multiple scattering.

We have also reported novel deep learning models for efficient analysis of multi-dimensional 3D data we acquire. In contrast to other work on image translation that demonstrated 2D prediction (34–36), our 2.5D architecture achieves 3D prediction with apparently similar or superior accuracy as the 3D prediction reported in (33) (Pearson correlation coefficient in 3D for nuclei prediction from brightfield images: 0.87 v.s. 0.7 reported in (33)), while being computationally more efficient. We note that Pearson correlation coefficient is affected by both the accuracy of the prediction as well as the noise in the target images. Thus a more direct comparison of model performances on the same dataset would be useful in the future. Also, 2.5D network can be applied to image data that only has a few z-slices without up- or down-sampling the data, making it useful for analysis of thin slices as well as 3D specimens. Even though we focus on image translation in this work, the same 2.5D network can be used for 3D segmentation. 3D segmentation using the 2.5D network bears additional advantages over 3D network because annotation can be done only on a subset of slices rather than the whole 3D volume, which would save significant amount of manual annotation time and efforts.

In comparison to Christiansen *et al.*’s 2D translation model (34) where the image translation was formulated as a pixel-wise classification task of 8-bit classes, our 2.5D translation model formulates the image translation as a regression task that allows prediction of much larger dynamic range of gray levels. While training a single model that predicts multiple structures seems appealing, this more complex task requires increasing the model size with the trade-off of longer model training time. Our modeling strategy to train one model to predict only one target allowed us to use significantly smaller models that can fit into the memory of a single GPU for faster training.

We systematically evaluated how the dimensions and input channels affect the prediction accuracy. Compared to previous work that predict fluorescence images from single label-free contrast (33–36), we show that higher prediction accuracy can be achieved by combining multiple label-free contrasts. Additionally, we demonstrated prediction of fluorescence images of tissue, while previous work has reported prediction of fluorescence images of cultured cells or bright-field images of histochemically-stained tissue (33–36).

A common shortfall of machine learning approaches is that they tend not to generalize well. We have shown that our data normalization and training process leads to models of myelination that generalize to two developmental time points in fig. 5. In contrast to reconstruction using physical models, we note that the errors or artifacts in the prediction by machine learning models are usually heterogeneous and highly dependant on the quality of training data and the input. Therefore, prediction errors made by machine learning models can be difficult to recognize in the absence of ground truth. For image translation in biological specimen with unknown structures, such as prenatal human brain, it would be crucial to also estimate the confidence interval of output values, which is an important area of research.

Several methods for tracing connectivity in the mouse brain at mesoscale have been developed (56), but they have not yet been scaled to human brain. The volume of fetal human brain during third trimester (10^5^mm^3^ −4×10^5^mm^3^) is 3 orders of magnitude larger than the volume of an adult mouse brain (~ 5 × 10^2^mm^3^). Our data shows that quantitative polarized light microscopy can provide label-free measurement of myelination and axon tract orientation at high density with ~ 1.5*µ*m resolution over centimeter spatial extents, in both primary human brain sections and mouse brain sections. Our results comparing human and mouse brain sections highlight the high degree of fasciculation of axonal bundles in the mouse brain compared with human. We anticipate that analysis of connectivity across large brain slices will be possible by adapting tractography algorithms developed for diffusion weighted-MRI measurements to our vectorial data. Scalable mapping of myelination and connectivity from archival tissue of postmortem brain may enable high-throughput mapping of mesoscale connectivity in large brains, leading to new insights into how information processing streams are organized across the developing and adult human brain.

## Conclusion

In summary, we report an integrative computational imaging approach that combines label-free imaging, reconstruction of physical properties using image formation models, and prediction of biological structures with data-driven deep neural network models. Our Stokes-formalism based reconstruction algorithms (https://github.com/mehta-lab/reconstruct-order) and computationally efficient U-Net variants (https://github.com/czbiohub/microdl) facilitate imaging and interpretation of label-free signatures across biological scales. Our approach enables simultaneous measurement of phase, retardance, orientation, and degree of polarization contrasts with diffraction-limited spatial resolution. These contrasts report variations in density, anisotropy, and scattering of the specimen. We demonstrated visualization of diverse biological structures: glomeruli and tubules in mouse kidney tissue, multiple organelles in cells, and axon tracts and myelination in mouse and human brain slices. We demonstrated multi-contrast 2.5D U-Net model for accurate and computationally efficient prediction of biological structures from label-free contrasts. Fluorescence predicted from label-free images is robust to inherent variability in labeling. We demonstrated accurate prediction of ordered F-actin and nuclei in heterogeneous tissue, as well as myelination in prenatal human brain tissue. We anticipate that our approach will enable scalable analysis of architectural order that underpins healthy and disease states of cells and tissues.

## Methods

### Model of image formation

We describe dependence of the polarization resolved images on the specimen properties using Stokes formalism (44, Ch.15). This representation allows us to accurately measure the polarization sensitive contrast at the image plane for every focal plane. First, we retrieve the coefficients of the specimen’s Mueller matrix that report linear birefringence, transmission, and depolarization. For brevity, we call them ‘Mueller coefficients’ of the specimen in this paper. Mueller coefficients are recovered from the polarization-resolved intensities and an instrument matrix that captures how Mueller coefficeints are related to intensities recorded by the microscope. Assuming that the specimen is mostly transparent, more specifically satisfies the first Born approximation (39), we reconstruct specimen phase, retardance, slow axis, and degree of polarization stacks from stacks of Mueller coefficients (fig. 1). The assumption of transparency is generally valid for the structures we are interested in, but does not necessarily hold when the specimen exhibits significant absorption or diattenuation. To ensure that the inverse computation is robust, we need to make judicious decisions about the light path, calibration procedure, and background estimation. A key advantage of Stokes instrument matrix approach is that it easily generalizes to other polarization diverse imaging methods - A polarized light microscope is represented directly by a calibrated instrument matrix.

For sensitive detection of birefringence, it is advantageous to suppress isotropic background by illuminating the specimen with elliptically polarized light and image with circular state of opposite handedness (25). For experiments reported in this paper, we acquired data by illuminating the specimen sequentially with right-handed circular and elliptical states and analyzed the transmitted light with left-handed circular state.

### Forward model: specimen properties → Mueller coefficients

We assume a weakly scattering specimen modeled by properties of net retardance *ρ*, orientation of the slow axis *ω*, transmission *t*, and depolarization *p*. The Mueller matrix of the specimen can be expressed as a product of two Mueller matrices, **M**_*t*_, representing the transmission and depolarization parts, and **M**_*r*_, representing the birefringent part of the specimen. The expression of **M**_*r*_ is a standard Mueller matrix of a linear retarder that can be found in (44, Ch.14), and **M**_*t*_ is expressed as

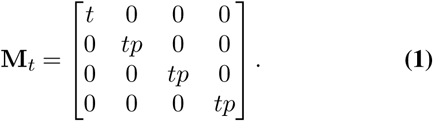

With **M**_*t*_ and **M**_*r*_, the Mueller matrix of the specimen is then given by

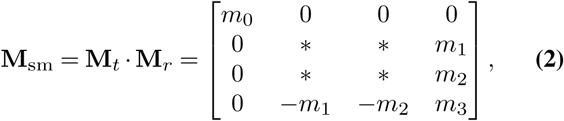

where * signs denote irrelevant entries that cannot be retrieved under our experiment scheme. The relevant entries that are retrievable can be expressed as a vector of Mueller coefficients, which is

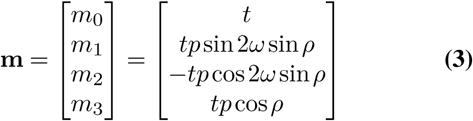

This vector is coincidentally the Stokes vector when right-handed circularly polarized light passing through the specimen. The aim of the measurement we describe in the following paragraphs is to accurately measure these Mueller coefficients at each point in the image plane of the microscope by illuminating the specimen and detecting the scattered light with mutually independent polarization states. Once a map of these Mueller coefficients has been acquired with high accuracy, the specimen properties can be retrieved from the above set of equations.

### Forward model: Mueller coefficients → intensities

To acquire the above Mueller coefficients, we illuminate the specimen with a series of right-handed circularly and elliptically polarized light (25). The Stokes vectors of our sequential illumination states are given by,

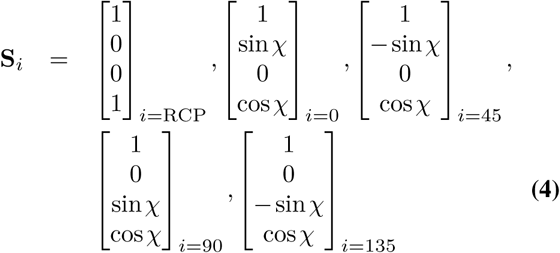

where *χ* is the compensatory retardance controlled by the LC that determines the ellipticity of the four elliptical polarization states.

After our controlled polarized illumination has passed through the specimen, we detect the light with the left-handed circular state by having a left-handed circular analyzer in front of our sensor. We express the Stokes vector before the sensor as

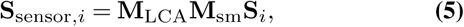

where *i* = {RCP, 0, 45, 90, 135} depending on the illumination states, and **M**_LCA_ is the Muller matrix of a left-handed circular analyzer (44, Ch.14). The detected intensity images are the first component of Stokes vector at the sensor under different illuminations (*I*_*i*_ = [**S**_sensor*,i*_]_0_). Stacking the measured intensity images to form a vector

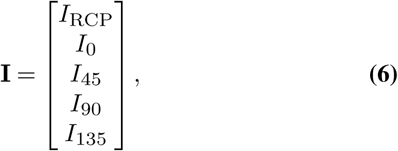

we can link the relationship between the measured intensity and the specimen vector through an ‘instrument matrix’ **A** as

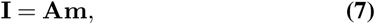

where

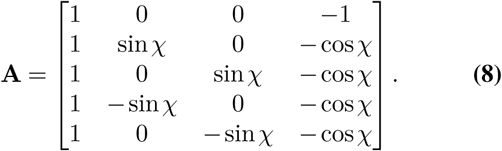

Each row of the instrument matrix is determined by the interaction between various illumination polarization states and the specimen’s properties. Any polarization-resolved measurement scheme can be characterized by an instrument matrix that transforms specimen’s polarization property to the measured intensities. Calibration of the polarization imaging system is then done through calibrating this instrument matrix.

### Computation of Mueller coefficients at image plane

Once the instrument matrix has been experimentally calibrated, the Stokes vector can be obtained from recorded intensities using its inverse (compare Eq. 7),

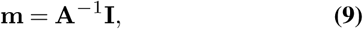

### Computation of background corrected specimen properties

We retrieved the vector of Mueller coefficients, **m**, by solving Eq. 9. Slight strain or misalignment in the optical components or the specimen chamber can lead to background that masks out contrast from the specimen. The background typically varies slowly across the field of view and can introduce spurious correlations in the measurement. It is crucial to correct the vector of Mueller coefficients for non-uniform back ground birefringence that was not accounted for by the calibration process. To correct the non-uniform background birefringence, we acquired background polarization images at the empty region of the specimen. We then transformed specimen (*i* = sm) and background (*i* = bg) vectors of Mueller coefficients as follows,

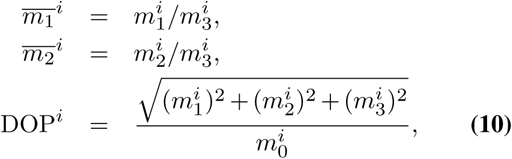

We then reconstructed the background corrected properties of the specimen: brightfield (BF), retardance (*ρ*), slow axis (*ω*), and degree of polarization (DOP) from the transformed specimen and background vectors of Mueller coefficients 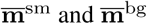 using the following equations:

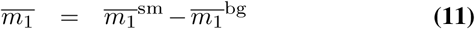

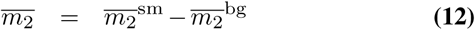

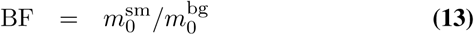

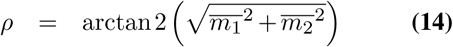

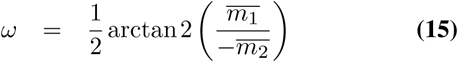

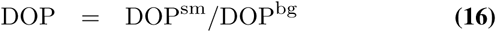

When the background cannot be completely removed using the above background correction strategy with a single background measurement, (i.e. the specimen has spatially varying background birefringence), we applied a second round of background correction on the measurements.

In this second round, we estimated the residual transformed background Mueller coefficients by fitting a low-order 2D polynomial surface to the transformed specimen Mueller coefficients. Specifically, we downsampled each 2048 × 2048 image to 64 × 64 image with 32 × 32 binning. We took the median of each 32 × 32 bin to be each pixel value in the downsampled image. We then fitted a second-order 2D polynomial surface to the downsampled image of each transformed specimen Mueller coefficient to estimate the residual background. With this newly estimated background, we performed another background correction. The effect of two rounds of the background corrections are shown in fig. 2-supplement S4.

### Phase reconstruction

As seen from Eq. 3, the first component in the vector of Mueller coefficients, *m*_0_, is equal to the total transmitted intensity of electric field in the focal plane. Assuming a specimen with weak absorption, the intensity variations in a Z-stack encode the phase information via the transport of intensity (TIE) equation (7). In the following, we leverage weak object transfer function (WOTF) formalism (17–22) to retrieve 2D and 3D phase from this TIE phase contrast and describe the corresponding inverse algorithm.

### Forward model for phase reconstruction

The linear relationship between the 3D phase and the through focus brightfield intensity was established in (17) with Born approximation and weak object approximation. In our context, we reformulated as

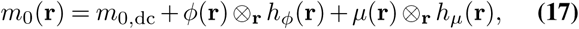

where **r** = (**r**_⊥_, z) = (*x, y, z*) is the 3D spatial coordinate vector, *m*_0,dc_ is the constant background of *m*_0_ component, ⊗_r_ denotes convolution operation over **r** coordinate, *φ* refers to phase, *µ* refers to absorption, *h_φ_*(**r**) is the phase point spread function (PSF), and *h_µ_*(**r**) is the absorption PSF. Strictly, *φ* and *µ* are the real and imaginary part of the scattering potential scaled by ∆*z/*2*k*, where ∆*z* is the axial pixel size of the experiment and *k* is the wavenumber of the incident light. When the refractive index of the specimen and that of the environment are close, the real and imaginary scaled scattering potential reduce to two real quantity, phase and absorption.

When specimen’s thickness is larger than the depth of field of the microscope (usually in experiments with high NA objective), the brightfield intensity stack contains 3D information of specimen’s phase and absorption. Without making more assumptions or taking more data, this problem is ill-posed because we are solving two unknowns from one measurement. Assuming the absorption of the specimen is negligible (18, 21, 22), which generally applies to transparent biological specimens, we turn this problem into a linear deconvolution problem, where 3D phase is retrieved.

When specimen’s thickness is smaller than the depth of field of the microscope (usually in experiments with low NA objective), the whole 3D intensity stack is coming from merely one effective 2D absorption and phase layer of specispecimen. We rewrite Eq. 17 as

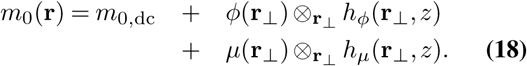

In this situation, we have multiple 2D defocused measurements to solve for one layer of 2D absorption and phase of the specimen.

### Inverse problem for phase reconstruction

With the linear relationship between the first component of the Mueller coefficients vector and the phase, we then formulated the inverse problem to retrieve 2D and 3D phase of the specimen.

When we recognize the specimen as a 3D specimen, we then use Eq. 17 and drop the absorption term to estimate specimen’s 3D phase through the following optimization algorithm:

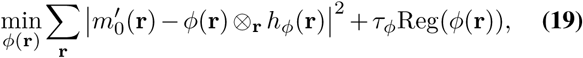

where 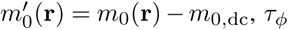 is the regularization parameter for applying different degree of denoising effect, and the regularization term depending on the choice of either Tikhonov or anisotropic total variation (TV) denoiser is expressed

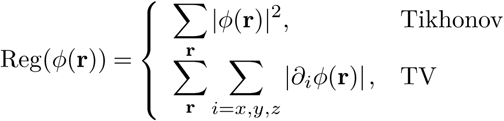

When using Tikhonov regularization, this optimization problem has an analytic solution that has previously described by (18, 21, 22). As for TV regularization, we adopted alternating minimization algorithm that is proposed and applied to phase imaging in (57) and (58), respectively, to solve the problem.

If we consider the specimen as a 2D specimen, we then turn Eq. 18 into the following optimization problem:

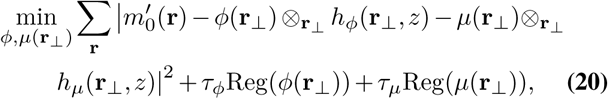

where we have an extra regularization parameter *τ_µ_* here for the absorption. When Tikhonov regularization is selected, the analytic solution similar to the one described in (59) is adopted.

When the signal to noise ratio of the brightfield stack is high, Tikhonov regularization gives satisfactory reconstruction in a single step with computation time proportional to the size of the image stack. However, when the noise is high, Tikhonov regularization can lead to high- to medium-frequency artifacts. Using iterative TV denoising algorithm, we can trade-off reconstruction speed with robustness to noise.

### Specimen preparation

Mouse kidney tissue slices (Thermo-Fisher Scientific) and mouse brain slices were mounted using coverglass and coverslip. U2OS cells were seeded and cultured in a chamber made of two strain-free coverslips that allowed for gas exchange. In the mouse kidney tissue slice, F-actin was labeled with Alexa Fluor 568 phalloidin and nuclei was labeled with DAPI.

Human prenatal brain samples were fixed with 4% paraformaldehye in phosphate-buffered solution (PBS) overnight, then rinsed with PBS, dehydrated in 30% sucrose/OCT compound (Agar Scientific) at 4C overnight, then frozen in OCT at −80 °C. Frozen samples were sectioned at 12 *µ*m and mounted on microscope slides. Sections were stained directly with red FluoroMyelin (Thermo-Fisher Scientific, 1:300 in PBS) for 20 minutes at room temperature, rinsed 3 times with PBS for 10 minutes each, then mounted with ProLong Gold antifade (Invitrogen) with a coverslip.

### Image acquisition and registration

We implemented LC-PolScope on a Leica DMi8 inverted microscope with Andor Dragonfly confocal for multiplexed acquisition of polarization-resolved images and fluorescence images. We automated the acquisition using Micro-Manager v1.4.22 and OpenPolScope plugin for Micro-Manager that controls liquid crystal universal polarizer (custom device from Meadowlark Optics, specifications available upon request).

We multiplexed the acquisition of label-free and fluorescence volumes. The volumes were registered using transformation matrices computed from similarly acquired multiplexed volumes of 3D matrix of rings from the ARGO-SIM test target (Argolight).

In transmitted light microscope, the resolution increases and image contrast decreases with increased numerical aperture of illumination. We used 63X 1.47 NA oil immersion objective (Leica) and 0.9 NA condenser to achieve a good balance between image contrast and resolution. The mouse kidney tissue slice was imaged using 100 ms exposure for 5 polarization channels, 200 ms exposure for 405 nm channel (nuclei) at 1.6 mW in the confocal mode, 100 ms exposure for 561 nm channel (F-actin) at 2.8 mW in the confocal mode. The mouse brain slice were also imaged using 30 ms exposure for 5 polarization channels. U2OS cells were imaged using 50 ms exposure for 5 polarization channels. For training the neural network, we acquired 160 non-overlapping 2048×2048×45 z-stacks of the mouse kidney tissue slice with Nyquist sampled voxel size 103nm×103nm×250nm. Human brain sections were imaged with a 10X objective and 0.2 NA condenser with a 200 ms exposure for polarization channels, 250 ms exposure for 568 channel (FluoroMyelin) in the epifluorescence mode. The full brain sections were imaged, approximately 200 images depending on the size of the section, with 5 z positions at each location.

### Data preprocessing for model training

The images were flat-field corrected. For training 3D models, the image volumes were upsampled along Z to match the pixel size in XY using linear interpolation. The images were tiled into 256×256 patches with a 50% overlap between patches for 2D and 2.5D models. The volumes were tiled into 128×128×96 patches for 3D models with a 25% overlap along XYZ. Tiles that had sufficient fluorescence foreground (2D and 2.5D: 20%, 3D: 50%) were used for training. Foreground masks were computed by summing the binary images of nuclei and F-actin obtained from Otsu thresholding in the case of mouse kidney tissue sections, and binary images of FluoroMyelin for the human brain sections.

Proper data normalization is essential for predicting the intensity correctly across different fields-of-views. We found the common normalization scheme where each image is normalized by its mean and standard deviation does not produce correct intensity prediction (fig. 5 - figure supplement S3). We normalized the images on the per dataset basis to correct the batch variation in the staining and imaging process across different datasets. To balance contributions from different channels during training of multi-contrast models, each channel needs to be scaled to similar range. Specifically, for each channel, we subtracted its median and divided by its inter-quartile range (range defined by 25% and 75% quantiles). We used inter-quartile range to normalize the channel because standard deviation underestimates the spread out of the distribution of highly correlated data such as pixels in images.

### Neural network architecture

We experimented with 2D, 2.5D and 3D versions of U-Net models fig. 3-supplement S1. Across the three U-Net variants, each convolution block in the encoding path consists of two repeats of three layers: a convolution layer, ReLU non-linearity activation, and a batch normalization layer. We added a residual connection from the input of the block to the output of the block to facilitate faster convergence of the model (47, 48). 2×2 downsampling is applied with 2×2 convolution with stride 2 at the end the each encoding block. On the decoding path, the feature maps were passed through similar convolution blocks, followed by upsampling using bilinear interpolation. Feature maps output by every level of encoding path were concatenated to feature maps in the decoding path at corresponding levels. The final output block had a convolution layer only.

The encoding path of our 2D and 2.5D U-Net consists of five layers with 16, 32, 64, 128 and 256 filters respectively, while the 3D U-Net consists of four layers with 16, 32, 64 and 128 filters each due to its higher memory requirement. The 2D and 3D versions use convolution filters of size of 3×3 and 3×3×3 with a stride of 1 for feature extraction and with a stride of 2 for downsampling between convolution blocks.

The 2.5D U-Net has the similar architecture as the 2D U-Net with following differences:

1. The 3D features maps are converted into 2D using skip connections that consist of a N×1×1 valid convolution.
2. Convolution filters in the encoding path are N×3×3, where N = 3,5,7 is the number of slices in the input.
3. In the encoding path, the feature maps are downsampled across blocks using N*×*2*×*2 average pooling.
4. In the decoding path, the feature maps were upsampled using bilinear interpolation by a factor of 1×2×2 and the convolution filters in the decoding path are of shape 1*×*3*×*3.

The 2D, 2.5D, 3D network with single channel input consisted of 2.0 M, 4.8M, 1.5M learnable parameters, respectively.

### Model training and inference

We randomly split the images in groups of 70%, 15%, and 15% for training, validation and test. The split are kept consistent across all model training to make the results comparable. All models are trained with Adam optimizer, L1 loss function, and a cyclic learning rate scheduler with a min and max learning rate of 5 × 10^*−*5^ and 6 × 10^*−*3^ respectively. The 2D, 2.5D, 3D network were trained on mini-batches of size 64, 16, and 4 to accommodate the memory requirements of each 3D model. Models were trained until the validation loss does not decrease for 20 epochs. The model with minimal validation loss was saved. Single channel 2D models converged in 6 hours, 2.5D model converged in 47 hours and the 3D model converged in 76 hours on NVIDIA Tesla V100 GPU with 32GB RAM.

As the models are fully convolutional, model predictions were obtained using full XY images as input for the 2D and 2.5D versions. Due to memory requirements of the 3D model, the test volumes were tiled along x and y while retaining the entire z extent (patch size: 96×512×512) with an overlap of 32 pixels along X and Y. The predictions were stitched together by linear blending the model predictions in the overlapping regions.

### Model evaluation

Pearson correlation and structural similarity index (SSIM) along the XY, XZ and XYZ dimensions of the test volumes were used for evaluating model performance.

The Pearson correlation coefficient between a target image *T* and a prediction image *P* is defined as

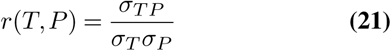

where *σ*_*TP*_ is the covariance of *T* and *P*, and *σ*_*T*_ and *σ*_*P*_ are the standard deviations of *T* and *P* respectively.

SSIM compares two images using a sliding window approach, with window size *N×N* (*N×N×N* for XYZ). Assuming a target window *t* and a prediction window *p*,

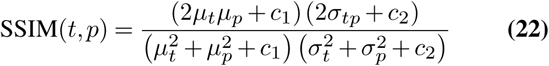

where *c*_1_ = (0.01*L*)^2^ and *c*_2_ = (0.03*L*)^2^, and *L* is the dynamic range of pixel values. Mean and variance are represented by *µ* and *σ*^2^ respectively, and the covariance between *t* and *p* is denoted *σ_tp_*. We use *N* = 7. The total SSIM score is the mean score calculated across all windows, SSIM score is the mean score calculated across all windows, SSIM 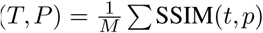 for a total of M windows. For XY and XZ dimensions, we compute one test metric per plane and for XYZ dimension, we compute one test metric per volume.

Importantly, it is essential to scale the the model prediction back to the original range before normalization for correct calculation of target-prediction SSIM. This is because unlike Pearson correlation coefficient, SSIM is not a scale-independent metrics.

## ACKNOWLEDGEMENTS

We thank Spyros Dermanis (CZ Biohub), Bing Wu (CZ Biohub), May Han (Stanford), and Ezzat Hashemi (Stanford) for providing the mouse brain slice used for acquiring data shown in Fig. 1. We thank Greg Huber, Loic Royer, Joshua Batson, Jim Karkanias, Joe DeRisi, and Steve Quake from the Chan Zuckerberg Biohub for numerous discussions. We also thank Eva Dyer from Georgia Tech for discussions about applications of the 2.5D models. This research was supported by the Chan Zuckerberg Biohub.

## Footnotes

1 To the first approximation, optical path length and retardance measured by the microscope are refractive index and birefringence of the specimen, respectively, integrated over the coherently illuminated volume of the specimen (6).

2 Slow axis is the orientation along which the anisotropic structure is denser. Higher density leads to slower propagation of light when the polarization of the light is oriented along the slow axis.

## Supplementary figures

**Figure 2 – supplement S1.**
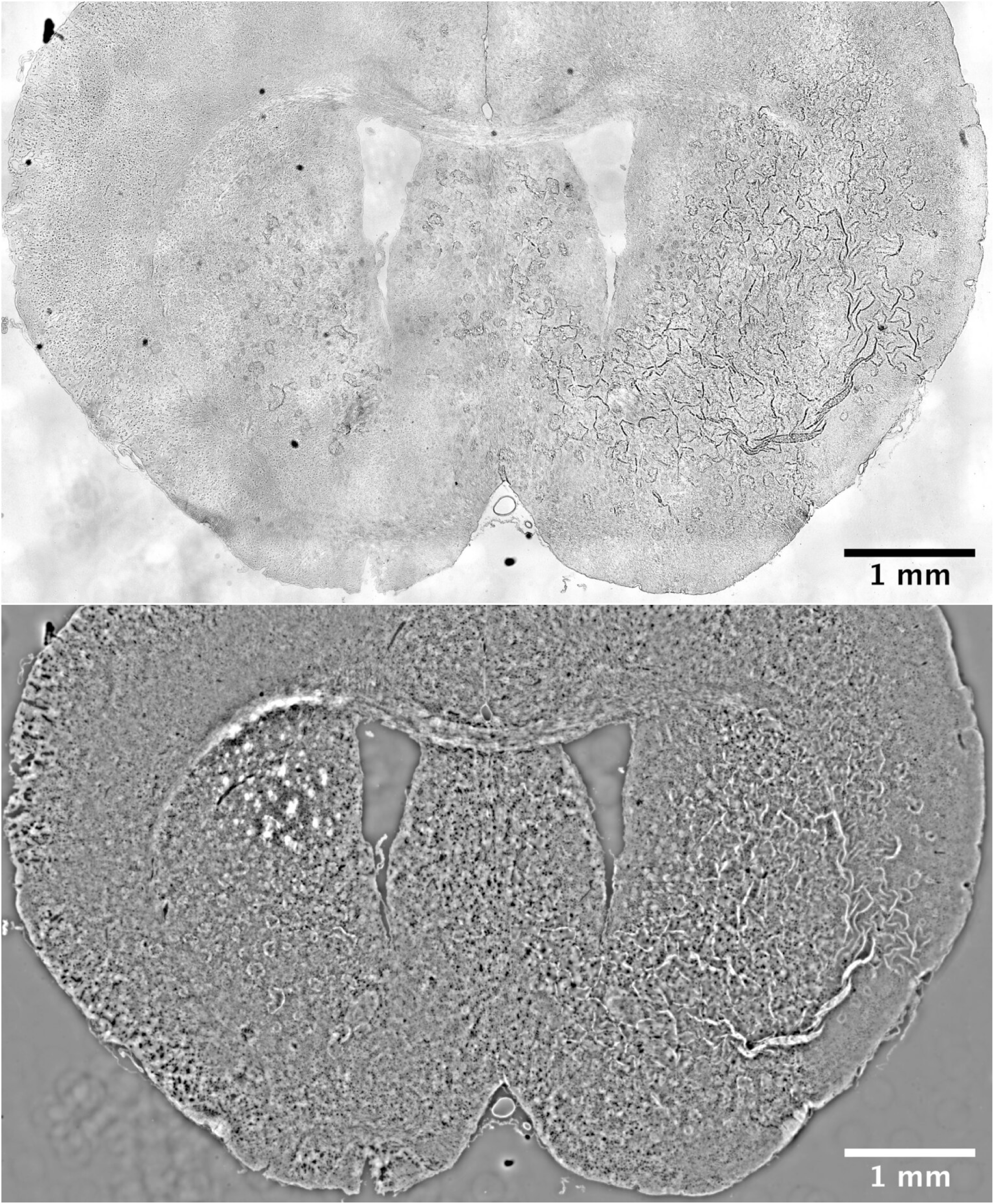
Comparison of background-corrected brightfield image (top) with quantitative phase image (bottom) of a mouse brain slice. The phase image reports density variations at higher contrast. These images are stitches of 48 fields of view and are substantially downsampled to reduce size.

**Figure 2 – supplement S2.**
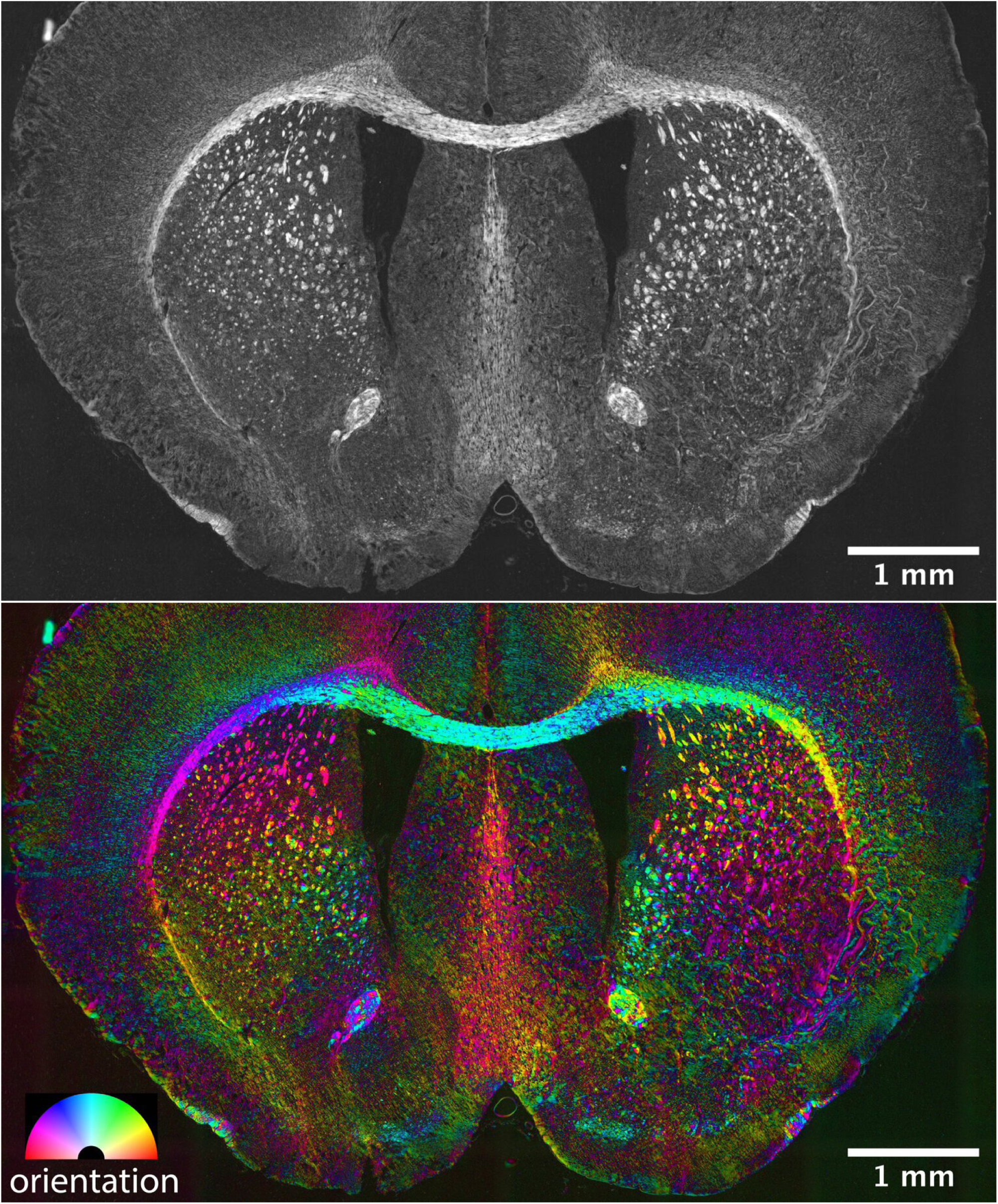
Retardance (top) and orientation (bottom) measurements of a mouse brain slice, which report structural anisotropy and slow axis orientation, respectively. We needed to compress the measured dynamic range of retardance and orientation by using gamma correction (0.5) to visualize less anisotropic gray matter in the presence of highly anisotropic white matter. These images are stitches of 48 fields of view and are substantially downsampled to reduce size. The peak retardance is 50nm.

**Figure 2 – supplement S3.**
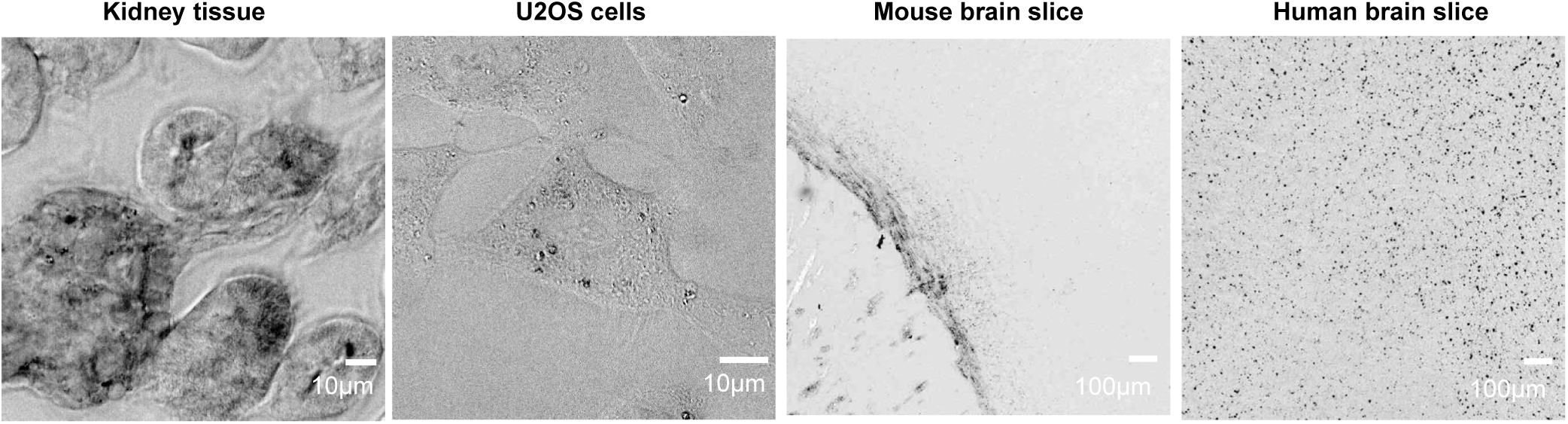
Degree of polarization (DOP) images for four specimens shown in fig. 2. When the specimen does not exhibit multiple scattering or diattenuation, DOP = 1. The DOP decreases when the specimen exhibits multiple scattering and increases when the diattenutation, i.e., polarization-dependent absorption.

**Figure 2 – supplement S4.**
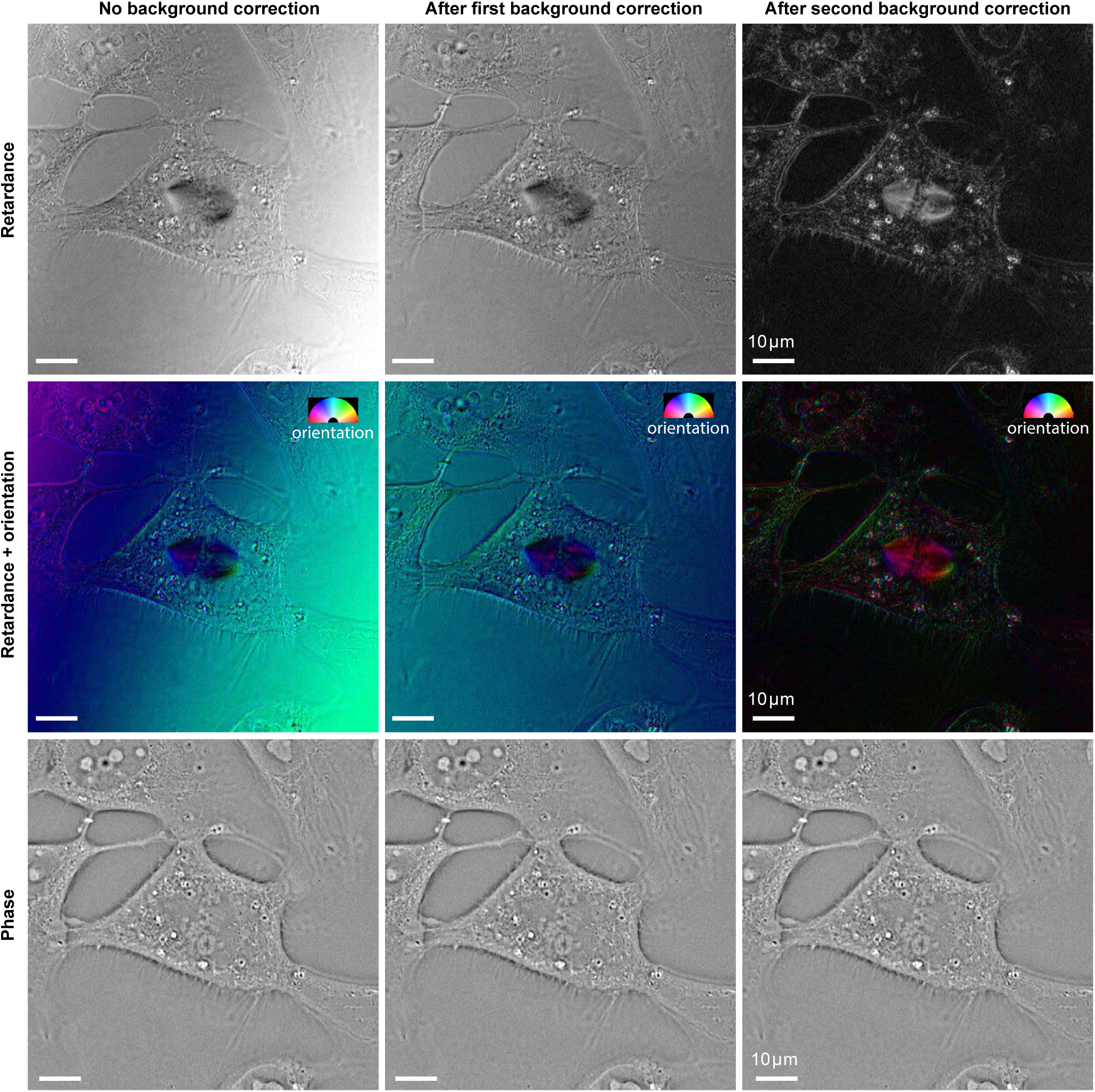
Effect of background correction methods on reconstructed retardance and phase of the U2OS cell. When the specimen has intrinsically low birefringence, background correction methods have a large impact on the reconstructed retardance and slow axis orientation. However, the background correction has no significant impact on phase reconstruction. (Left column) Reconstructions without background correction. (Middle column) Background-corrected reconstruction using an experimental images of empty region next to the cells (Right column) Background-corrected reconstruction using images estimated by fitting a very smooth surface to specimen image.

**Figure 3 – supplement S1.**
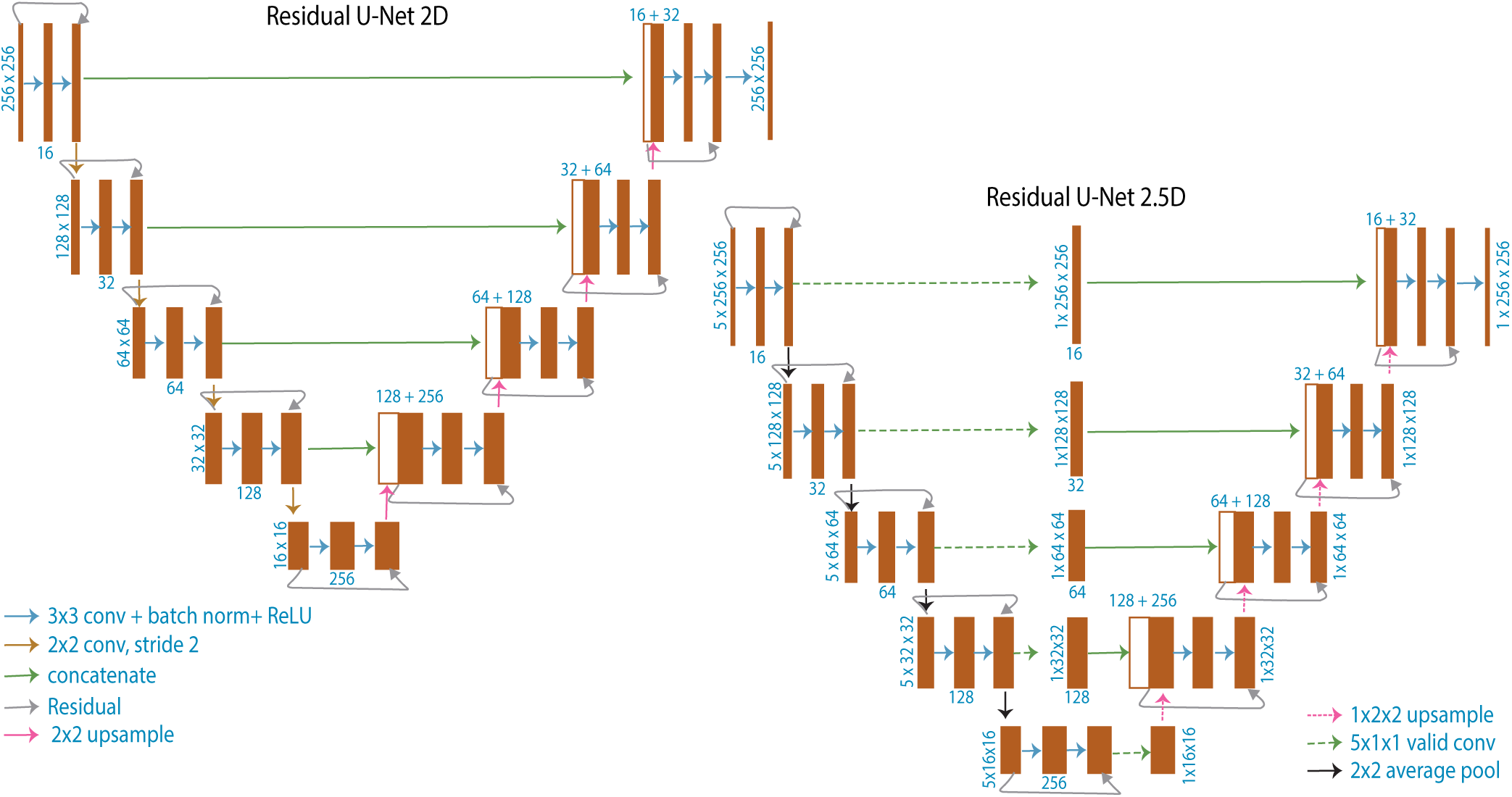
Schematic illustrating U-Net architectures: Schematic of 2D U-Net model used for translating slice→slice and 2.5D U-Net model used for translating stack→slice. The 3D U-Net model used for translating stack→stack is similar to the 2D U-Net, but uses 3D convolutions instead of 2D and is 4 layers deep instead of 5 layers deep.

**Figure 5 – supplement S1.**
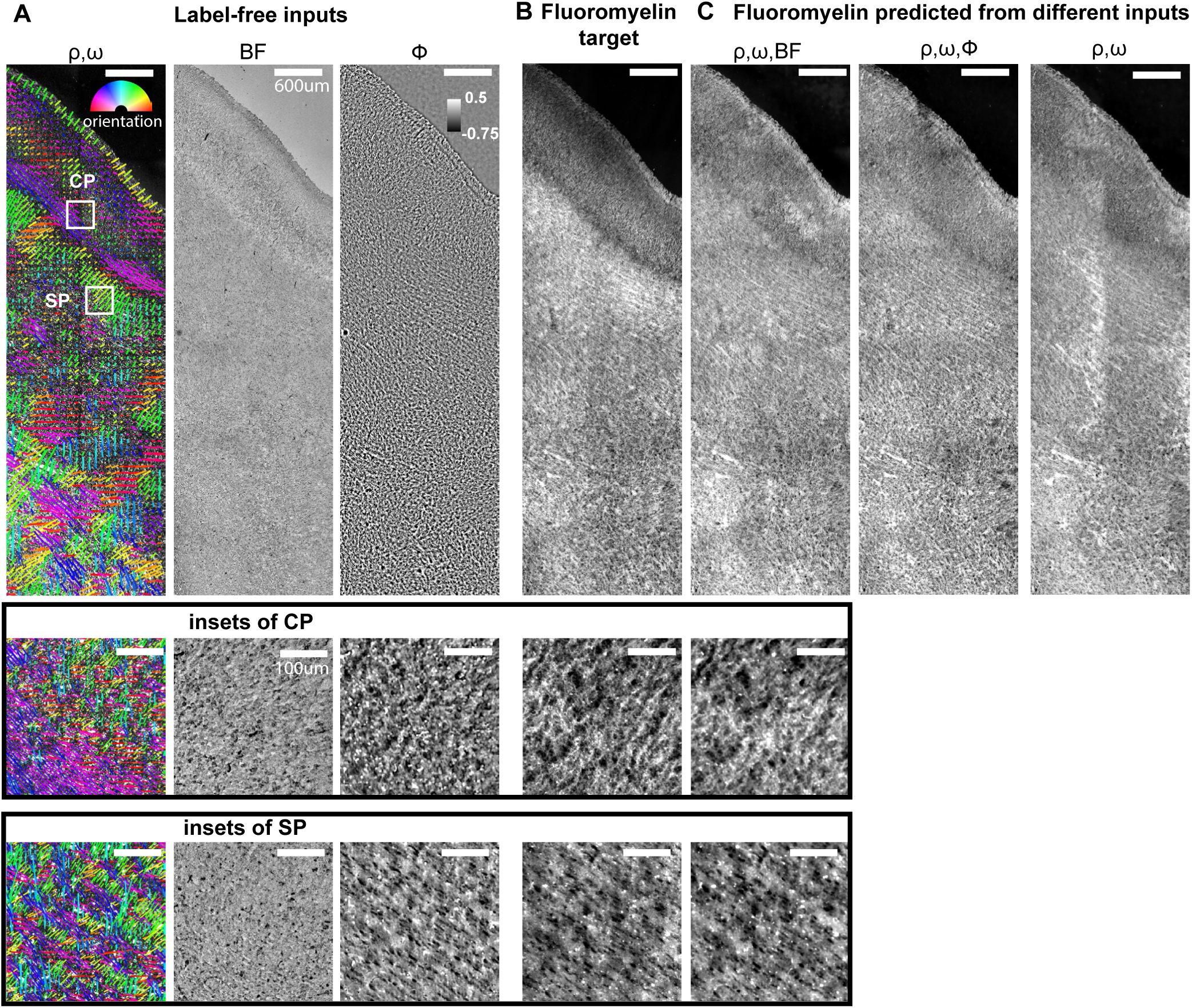
Label-free images and predicted Fluoromyelin images of a slice of human brain at gestational week 24 (GW24): (A) retardance (*ρ*), orientation (*ω*), brightfield (BF), and phase (*φ*) images from a field of view drawn from a test dataset, (B) experimental Fluoromyelin (target), (C) Fluoromyelin predicted by image translation models using different label-free images as inputs: (left to right) retardance, orientation, and brightfield as input to 2.5D model; retardance, orientation, and phase as input of 2D model; retardance and orientation as input to 2.5D model. Insets at the bottom show label-free, target, and predicted images within cortical plate (CP) and subplate (SP) at higher magnification. Scale bars: (A-C) 600*µ*m, (insets) 100*µ*m.

**Figure 5 – supplement S2.**
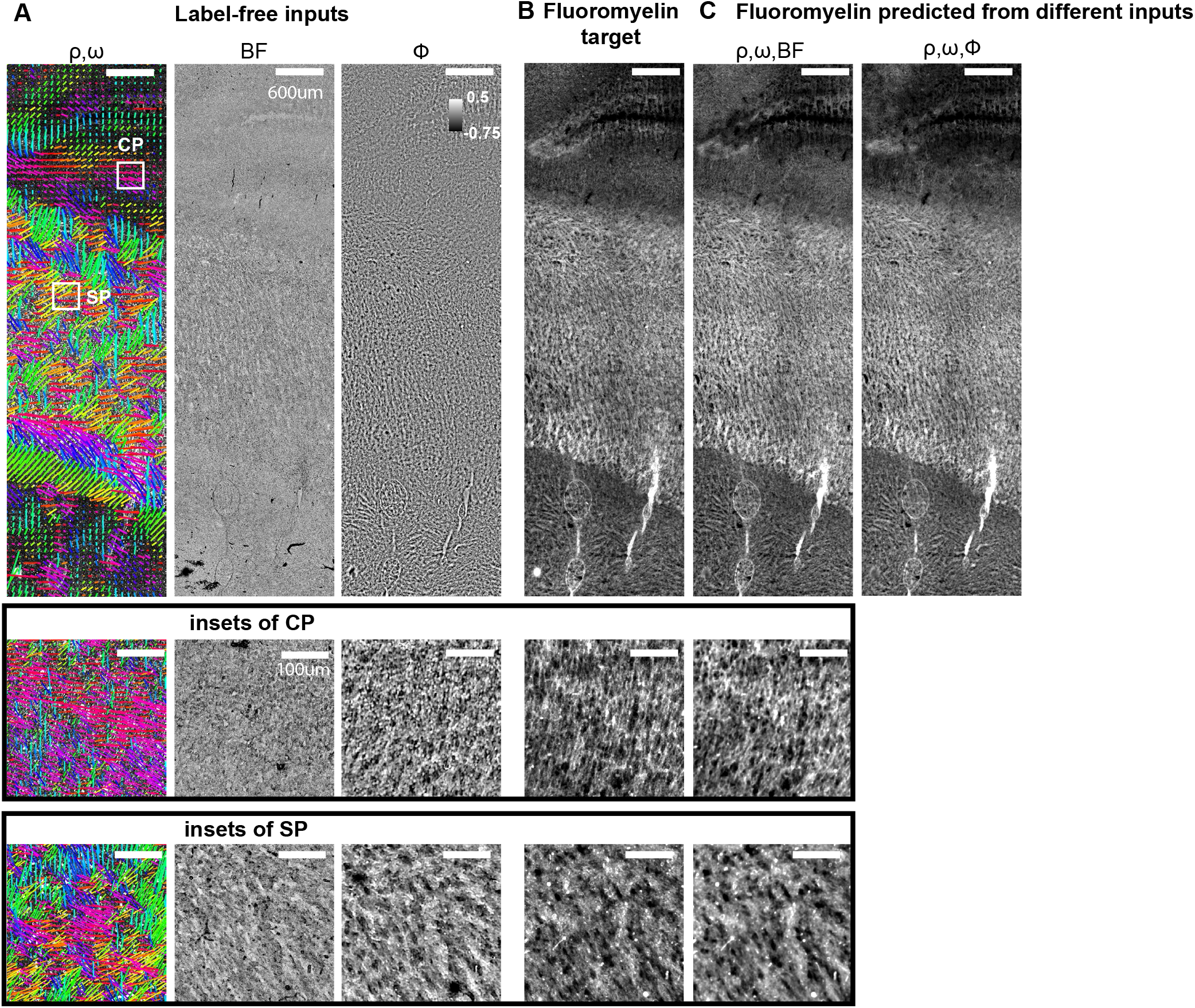
Label-free images and predicted Fluoromyelin images of a slice of human brain at gestational week 20 (GW20): The data is presented in the same way as fig. 5-supplement S1.

**Figure 5 – supplement S3.**
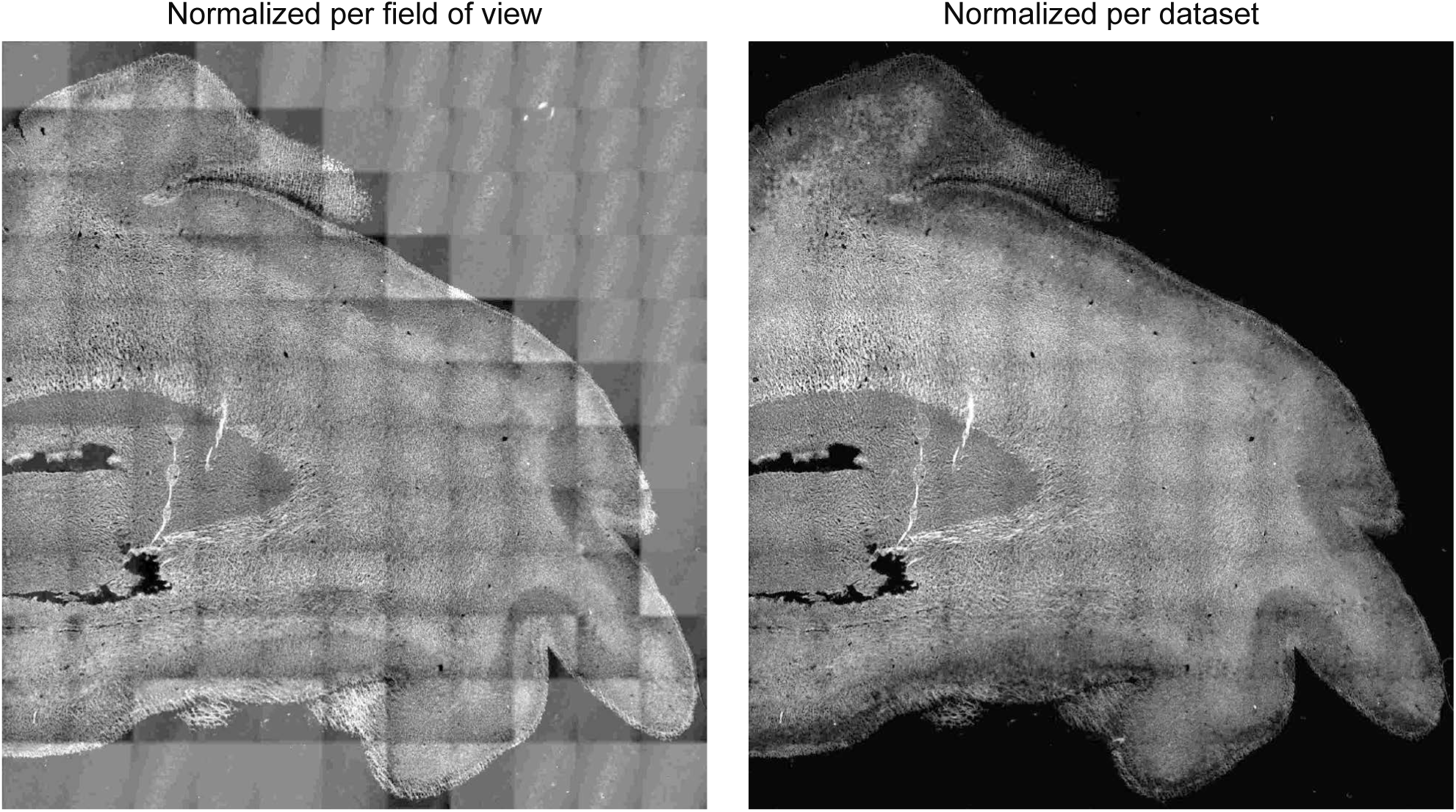
Normalizing training data per dataset yields prediction with correct dynamic range of intensity: Predictions of FluoroMyelin in GW20 human brain tissue slice with training data normalized per field of view (left) and across dataset (right)

## Videos

**Video 1.**
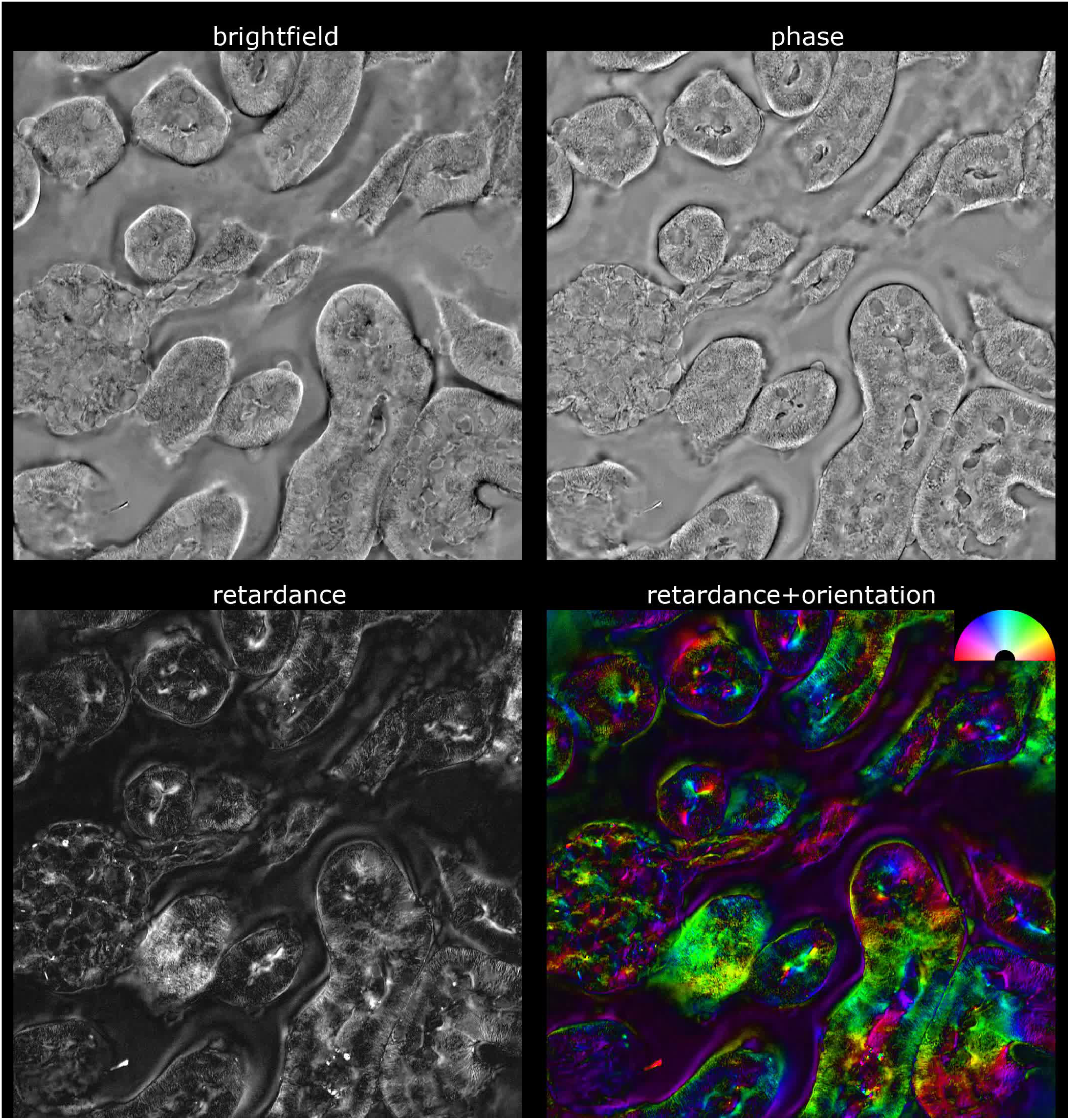
Z-stacks of brightfield, phase, retardance, and orientation images of mouse kidney tissue. The same field of view is shown in fig. 2, fig. 3, and fig. 4.

**Video 2.**
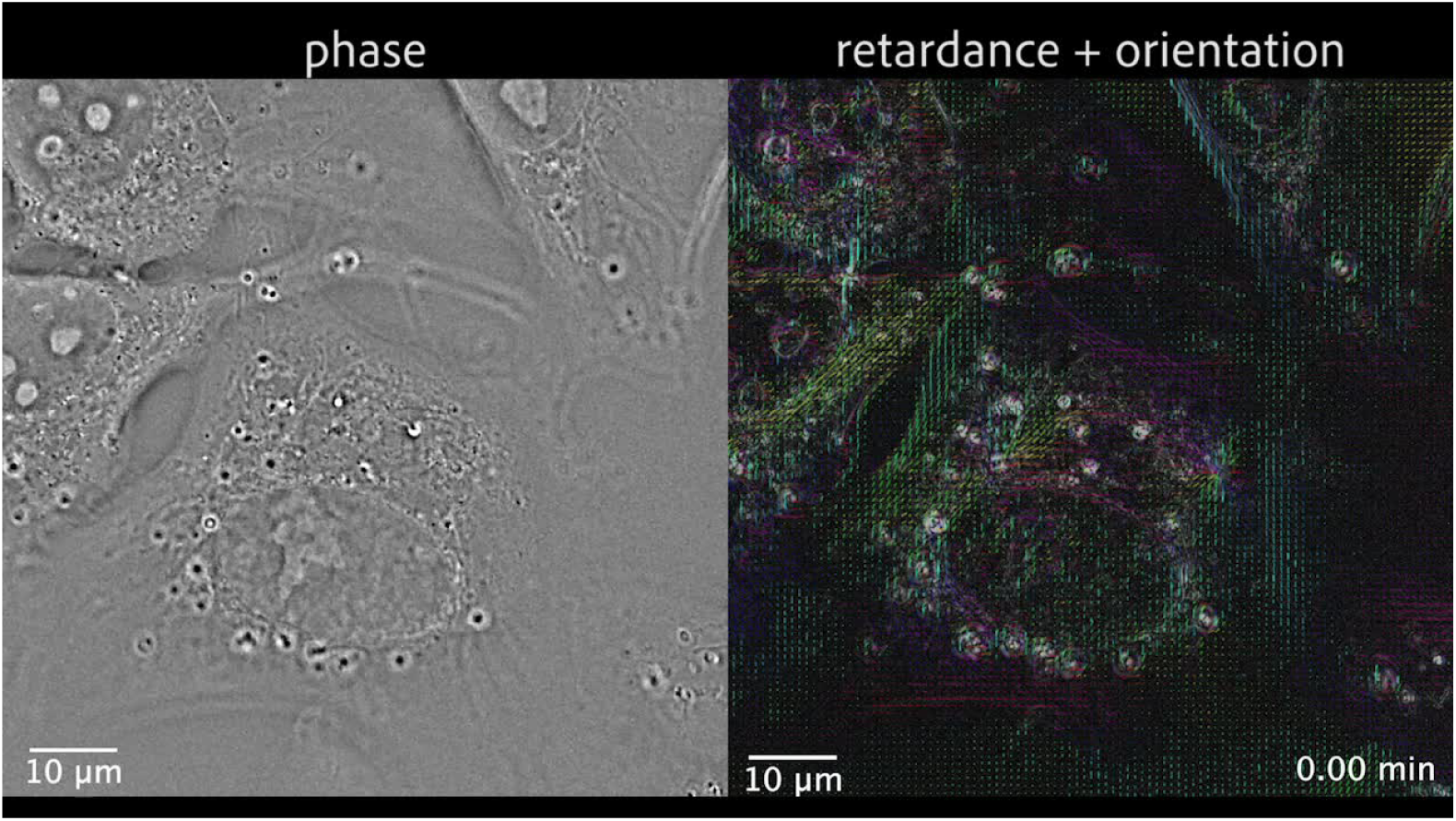
Time-lapse of phase, retardance, and slow axis orientation in a dividing U2OS cell shows differences in density and anisotropy of organelles. he same field of view is shown in fig. 2

**Video 3.**
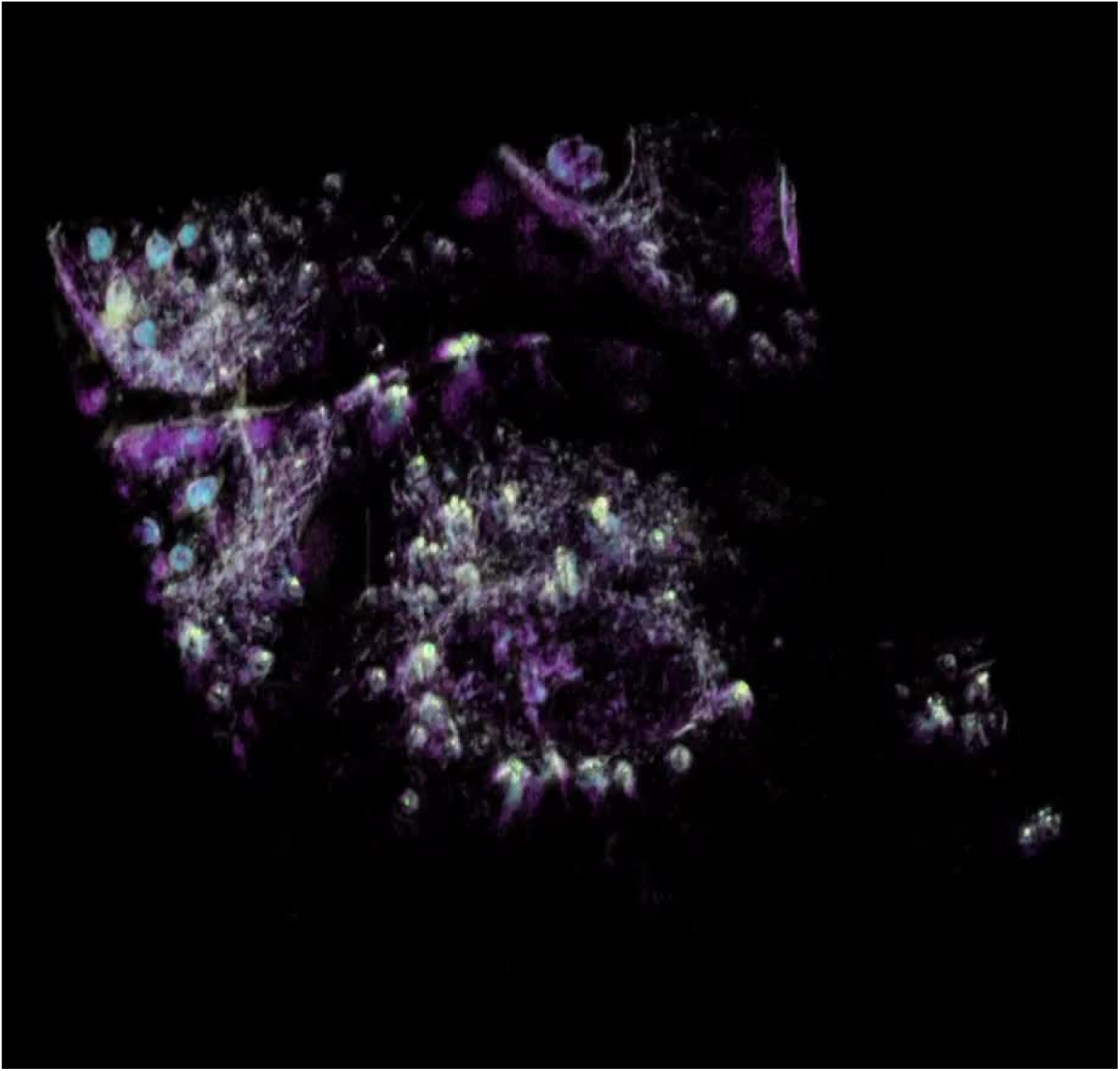
3D rendering of the time-lapse showing diverse structures color coded by their retardance and phase in U2OS cells shown in fig. 2. Phase values from low to high are color-coded with magenta, cyan, and green, respectively. Retardance values are color-coded with yellow. With this pseudo-color scheme, cytoplasm appears in magenta, chromatin appears in magenta and changes to blue as it condenses to chromosomes, nucleoli appear in blue, lipid vesicles appear in green surrounded by yellow rings, and spindles appear in yellow.

**Video 4.**
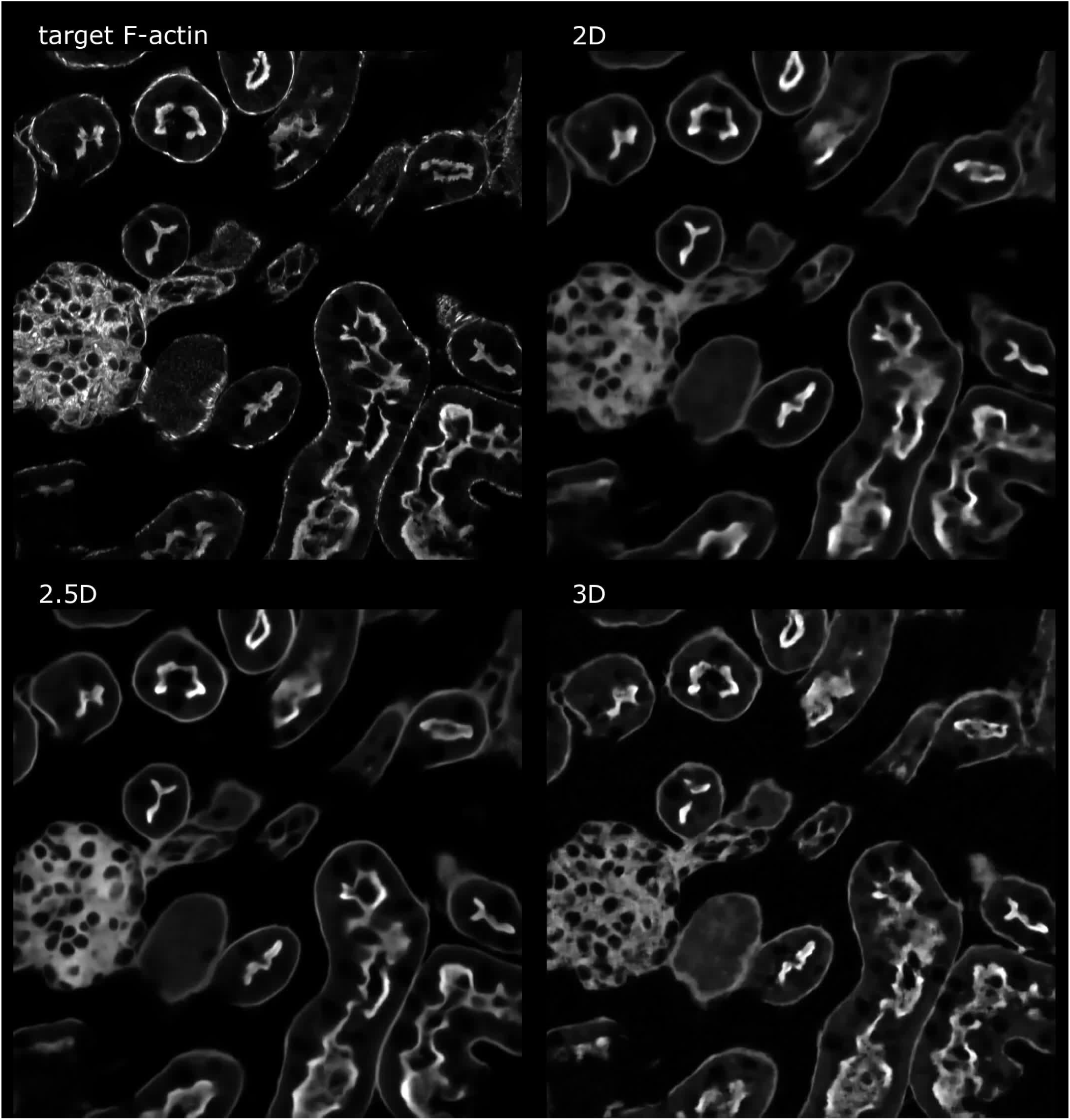
Through focus series showing 3D F-actin distribution in the test field of view shown in fig. 3: We show F-actin distribution (labeled with phalloidin-AF568) acquired on a confocal microscope (target), as well as predicted from 2D, 2.5D, and 3D models trained to translate retardance distribution into fluorescence distributions.

**Video 5.**
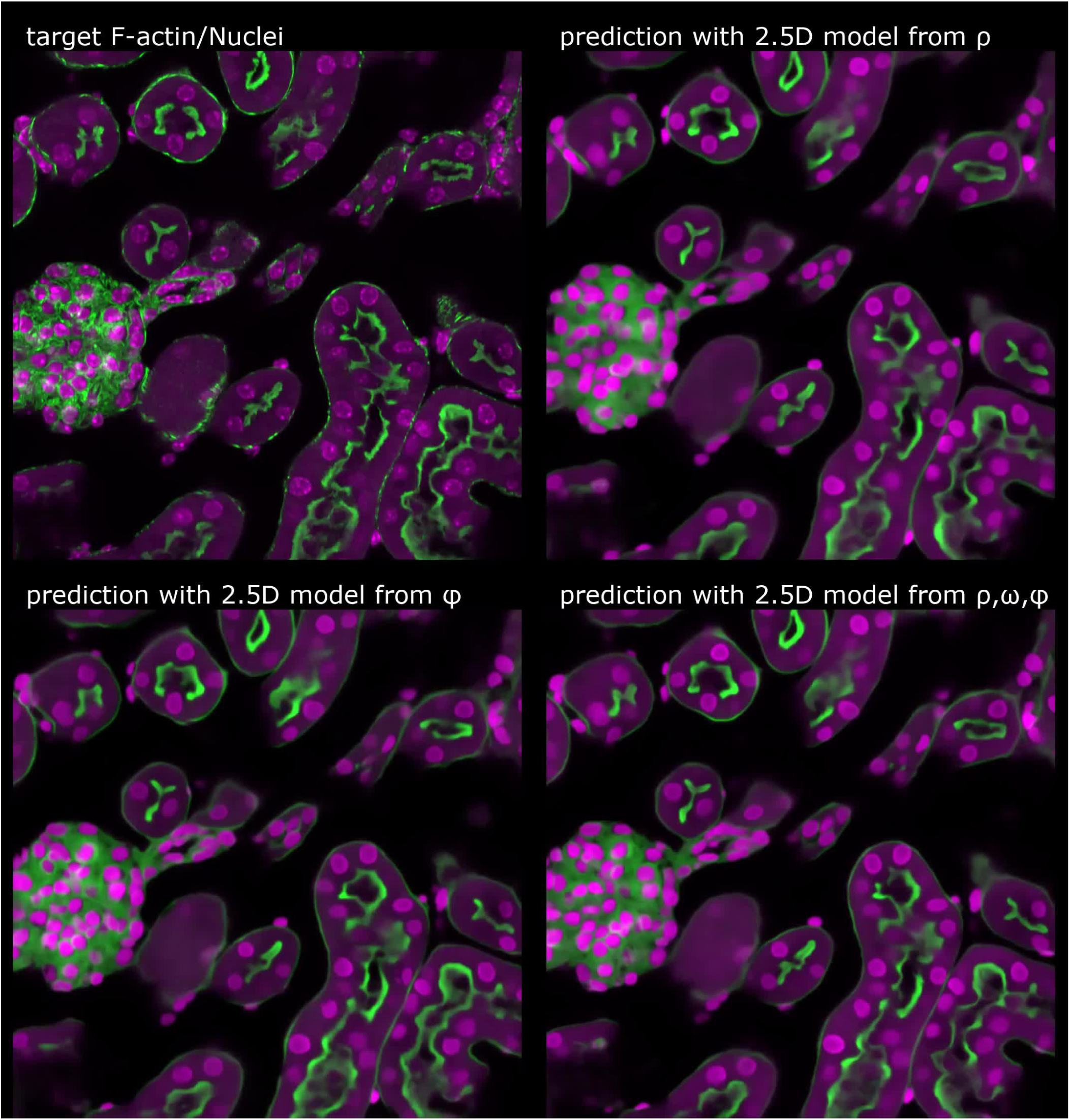
Through focus series showing 3D F-actin and nuclei distribution in the test field of view shown in fig. 4: We show overlays of F-actin (phalloidin-AF568) distribution in green and nuclei (DAPI) distribution in magenta DAPI as acquired on confocal (target) and as predicted from models. Predictions were obtained from 2.5D models trained on retardance (*ρ*) alone, phase (*φ*) alone, and on combination of retardance, orientation, phase (*ρ, ω, φ*).

## Notes

#### Summary of Updates

We have added supplementary figure (Fig. 2 - supplement 4) to illustrate background correction method and revised the text.

https://github.com/mehta-lab/reconstruct-order

https://github.com/czbiohub/microdl

